# Cyclic uniaxial mechanical load enhances chondrogenesis through entraining the molecular circadian clock

**DOI:** 10.1101/2021.10.26.465847

**Authors:** Judit Vágó, Éva Katona, Roland Takács, Klaudia Dócs, Tibor Hajdú, Patrik Kovács, Róza Zákány, Daan R. van der Veen, Csaba Matta

## Abstract

The biomechanical environment plays a key role in regulating cartilage formation, but current understanding of mechanotransduction pathways in chondrogenic cells is incomplete. Amongst the combination of external factors that control chondrogenesis are temporal cues that are governed by the cell-autonomous circadian clock. However, mechanical stimulation has not yet directly been proven to modulate chondrogenesis *via* entraining the circadian clock in chondroprogenitor cells. The purpose of this study was to establish whether mechanical stimuli entrain the core clock in chondrogenic cells, and whether augmented chondrogenesis caused by mechanical loading was at least partially mediated by the synchronised, rhythmic expression of the core circadian clock genes, chondrogenic transcription factors, and cartilage matrix constituents at both transcript and protein levels. We report here, for the first time, that cyclic uniaxial mechanical load applied for 1 hour for a period of 6 days entrains the molecular clockwork in chondroprogenitor cells during chondrogenesis in limb bud-derived micromass cultures. In addition to the several core clock genes and proteins, the chondrogenic markers *SOX9* and *ACAN* also followed a robust sinusoidal rhythmic expression pattern. These rhythmic conditions significantly enhanced cartilage matrix production and upregulated marker gene expression. The observed chondrogenesis-promoting effect of the mechanical environment was at least partially attributable to its entraining effect on the molecular clockwork, as co-application of the small molecule clock modulator longdaysin attenuated the stimulatory effects of mechanical load. This study suggest that an optimal biomechanical environment enhances tissue homeostasis and histogenesis during chondrogenesis at least partially through entraining the molecular clockwork.

## 1. Introduction

Cartilage is a specialised connective tissue with major structural and mechanical roles in the body. Articular cartilage is involved in load bearing, providing shape, cushioning and lubricating diarthrodial joints ^1^. Given that chondrocytes only occupy 1–3% of the tissue volume in mature cartilage, the unique biomechanical properties of the tissue are attributable to the composition and organization of the extracellular matrix (ECM), which is a complex network of collagens, proteoglycans, other non-collagenous proteins, and constituent water ^2^. The special architecture of the ECM provides cartilage with unique biomechanical properties in terms of compression, shear, and tension.

During the complex process of chondrogenesis, chondroprogenitor mesenchymal cells differentiate into chondroblasts and then chondrocytes, which produce cartilage ECM in response to internal and/or external stimuli ^3^. These stimuli include a combination of chemical, biological, and physical cues in the stem cell niche of progenitor cells, the specific combination and timing of which are essential for chondrogenesis ^4^. One of the most important external factors relevant to articular cartilage is the biomechanical environment. External mechanical load of an appropriate magnitude and frequency is essential for maintenance of the mature articular cartilage phenotype ^5^. The stimulatory effect of biomechanical forces on ECM synthesis by chondrocytes has been well documented ^6^; dynamic loading of mature cartilage enhances the production of glycosaminoglycans, cartilage oligomeric matrix protein (COMP), and collagen type II ^7^. Regular moderate physical activity has been shown to be necessary to maintain the healthy composition of articular cartilage ECM ^8^.

Biomechanical stimulation also plays a key role in regulating cartilage growth during the development of the skeletal system ^6^. The importance of movement during embryonic chondrogenesis and joint formation has been demonstrated using paralysed chicken embryos ^6,9^. Dynamic compressive load has been reported to stimulate epiphyseal cartilage growth ^10^, and variations in mechanical loading of articular cartilage have been proposed to modulate cartilage thickness ^11^. Both static and uniaxial cyclic compression have been reported to enhance the chondrogenic differentiation of embryonic limb bud-derived mesenchymal cells through the upregulation of collagen type II, aggrecan core protein, and the chondrogenic transcription factor SOX9 ^12,13^. Our group has documented the stimulatory effect of dynamic uniaxial mechanical load on chick limb bud-derived micromass cultures *via* the PKA/CREB-SOX9 and PP2A pathways ^14^. However, despite the emerging results on the molecular mechanisms mediated by mechanical load, current understanding of the mechanotransduction pathways in chondrogenic cells is still generally lacking.

Amongst the combination of chemical, biological, and physical factors known to guide mesenchymal stem cell (MSC) differentiation, a temporal cue recently discovered to be involved in the process is governed by the circadian clock ^15,16^. The molecular mechanisms that regulate the circadian clock in somatic cells rely on a network of autoregulatory transcriptional-translational feedback loops (TTFL), which drives the rhythmic expression patterns of the clock genes at the core of the circadian clock ^17^. The primary TTFL is controlled by the transcription factors CLOCK and BMAL1 which, after forming a heterodimer, bind to regulatory elements within target core clock genes including period (*PER1, PER2*, and *PER3*) and cryptochrome (*CRY1* and *CRY2*), as well as many clock-controlled genes (CCGs). PER and CRY proteins in turn multimerise and inhibit CLOCK/BMAL1 activity, thus blocking their own transcription ^17^. The CLOCK:BMAL1 heterodimers also regulate the transcription of retinoic acid-related orphan nuclear receptors, REV-ERBs and RORs, which bind to retinoic acid-related orphan receptor response elements (ROREs) present in the *BMAL1* promoter. REV-ERBs repress transcription of *BMAL1*, whereas RORs activate *BMAL1* transcription ^17^.

The ∼24 hour period of the endogenous molecular clockwork is reset daily by external timing cues, known as *Zeitgebers*. Whilst exposure to direct sunlight is the primary factor in entraining the main circadian clock in the hypothalamus, peripheral tissues are not light sensitive in mammals. Peripheral clocks thus entrain to systemic cues driven by the central hypothalamic clock, such as growth factors and hormones (e.g., glucocorticoids), but various other cues in their local environment also have a well-established role as *Zeitgeber* for peripheral clocks ^17^. An emerging entrainment mechanism for the circadian clock is mechanical stimulation. Rhythmic mechanical loading of the stretch-sensitive chordotonal organs can synchronise the *Drosophila* circadian clock ^18^. The mechanical environment of the epithelial stem cell niche controls the amplitude of the oscillation of the molecular clock ^16^. Although the specific molecular mechanisms whereby mechanical load entrains the circadian clockwork are not fully explored, *CLOCK* has been found to be mechanosensitive in chondrocytes ^19^. Mechanical stress loading has been shown to affect the circadian rhythm in developing skeletal muscle through modulating core clock gene expression ^20^.

Being necessary for maintaining healthy cartilage ^21^, recent results indicate that the clock genes can directly influence MSC differentiation, including adipogenesis ^22^ and chondrogenesis ^15^.

Recently, interactions between biological clock proteins and developmental pathways in chondrogenesis have become an emerging area of research. Indian hedgehog (IHH), a central regulator of chondrocyte differentiation, and its receptor Ptch1, are directly controlled by the circadian clock in chondrocytes ^23,24^. Key transcription factors relevant to chondrogenesis including SOX6 and SOX9 are likely also controlled by the biological clock during chondrocyte differentiation ^15,25^. The expression of genes involved in cartilage ECM turnover including *ACAN, MMP13*, and *COL2A1* have been shown to oscillate over the course of a day in chondrocytes ^26^. However, cyclic mechanical stimulation has not yet been shown to enhance chondrogenesis *via* entraining the circadian clock in chondroprogenitor cells. Here we report for the first time that cyclic uniaxial mechanical load applied for 1 hour for a period of 6 days synchronises the molecular clockwork during chondrogenesis in limb bud-derived micromass cultures and may enhance chondrogenic differentiation *via* entraining the circadian rhythm.

## 2. Materials and Methods

### 2.1. Experimental setup

Micromass cell cultures undergoing spontaneous *in vitro* chondrogenic differentiation were set up using embryonic chicken limb bud-derived chondroprogenitor cells. Cultures were subjected to 60 min of uniaxial dynamic loading every 24 hours for 6 days using our custom-made bioreactor ^14^. The start of the last mechanical loading regime on culturing day 6 was considered time point 0. Samples were then collected every 8 hours between 24-and 72-hours post-stimulation for evaluating the possible rhythmicity in the transcript and protein level expression of the core clock components and chondrogenic markers. Control, unstimulated cultures from the same biological replicates received a medium change with fresh DMEM containing 10% FBS at time point 0 on culturing day 6, and were harvested at the same time points as the cultures receiving mechanical load. The transcript expression patterns of clock genes (*BMAL1, PER2, PER3, CRY1, CRY2, REV-ERB*) and osteo-chondrogenic marker genes (*SOX6, SOX9, RUNX2, COL2A1, ACAN*) were studied using qRT-PCR and were analysed for circadian rhythms using cosine fitting. Time-dependent protein expression patterns of BMAL1, CRY1, PER3, RUNX2, SOX9, and aggrecan were analysed using Simple Western Wes immunoassay. The effects of the circadian clock regulator longdaysin on chondro-and osteogenesis, in combination with mechanical load, were evaluated using histological assessment following metachromatic staining, and by monitoring the expression levels of osteo-chondrogenic markers using qRT-PCR after the last mechanical stimulation on day 6. Analyses were performed on 3 biological replicates (*N* = 3). The workflow is summarised in *Fig. 1*.

**Figure 1.**
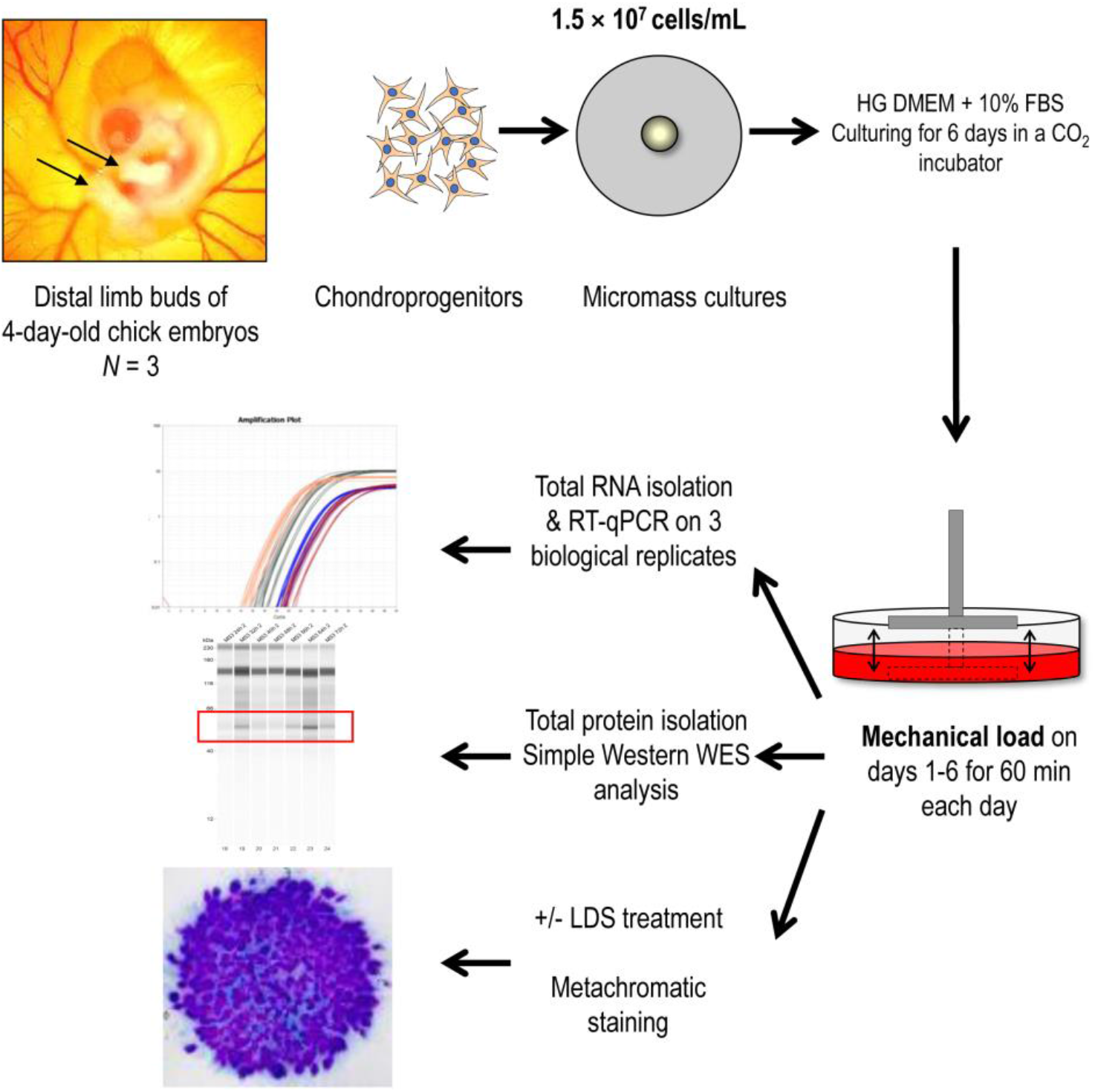
Experimental workflow. HG DMEM, high glucose-containing DMEM medium (4.5 g/L); LDS, longdaysin

### 2.2. Primary micromass cell cultures

Chondrifying micromass cell cultures were established from early-stage chicken embryos as previously described ^15^. To obtain a sufficiently high yield of primary chondrogenic cells, distal parts of the limb buds of ∼100 embryos were collected for each experiment. Briefly, distal parts of forelimbs and hindlimbs of embryos were isolated, pooled and dissociated in trypsin–EDTA (Merck, Darmstadt, Germany) for 1 hour. The dissociated limb buds were then filtered through a 20-μm pore size cell strainer (Merck) to generate a single-cell suspension of chondrogenic mesenchymal cells. Cells were then pelleted and resuspended in high glucose DMEM culture medium (Lonza, Basel, Switzerland) supplemented with 10% FBS (Lonza) at a concentration of 1.5 × 10^7^ cells/mL. 100 μL droplets were inoculated into 6-well plates (Eppendorf, Hamburg, Germany). After allowing the cells to adhere to the surface for 2 hours in a CO2 incubator (37°C, 5% CO2 and 90% humidity), 2 mL of DMEM supplemented with 10% FBS and 1% Penicillin/Streptomycin was added. The day of inoculation was considered as day 0 of culturing. Cultures were maintained at 37°C in a CO2 incubator. The medium was changed every second day, after mechanical load was applied. Experiments were performed in three biological replicates (*N* = 3).

### 2.3. Uniaxial dynamic mechanical load

Chondrifying micromass cultures grown in 6-well plates were subjected to uniaxial cyclic compressive force (approx. 0.6 kPa, 0.05 Hz) on every culturing day starting on day 1 for 60 min at exactly the same time of each day using a custom-made mechanical stimulator unit (for a detailed description of the bioreactor, please see ^14^). Mechanical load was carried out in normal DMEM culture medium containing 10% FBS. The set-up applies cyclic load transmitted *via* the culture medium to micromass cultures. The loading scheme comprises two main components: (a) uniaxial intermittent compressive force (hydrostatic pressure); and (b) fluid shear stress resulting from medium flow. The start of the last mechanical loading regime on culturing day 6 was considered time point 0. Samples were collected every 8 hours between 24-and 72-hours post-stimulation after day 6. Control, non-stimulated cultures from the same experiments received a medium change with fresh DMEM containing 10% FBS at time point 0, and harvested at the same time points as colonies receiving mechanical load. Harvested micromass cultures were stored at –80°C until RNA isolation.

### 2.4. Modulating the function of the molecular clock by longdaysin

Longdaysin (LDS; Cat. No.: SML0127; Merck) is a molecular clock modifier which blocks casein kinase I (CKI) and some other kinases ^27^. CKI phosphorylates the PER protein and promotes its degradation ^28^. LDS was dissolved in dimethyl sulfoxide (DMSO) at 5 mM and then administered to micromass cultures at the final concentration of 5 µM for the duration of the mechanical load on each culturing day. For these experiments, control cultures were treated with equal amount of the vehicle (DMSO).

### 2.5. RNA isolation and reverse transcription

Total RNA was isolated from micromass cultures using the TRI Reagent (Applied Biosystems, Thermo Fisher Scientific, Waltham, MA, USA) according to the instructions of the manufacturer. After the addition of 20% chloroform, samples were centrifuged at 4°C at 10,000×*g* for 15 min. Samples were incubated in 500 µL of RNase free isopropanol at –20°C for 1 hour, then total RNA was harvested in RNase-free water and stored at –80°C. RNA concentration and purity were determined by a Nanodrop 1000 UV-Vis spectrophotometer (Thermo Fisher Scientific). For gene expression analyses, 1 μg of RNA was reverse transcribed into complementary cDNA using the High-Capacity cDNA Reverse Transcription Kit (Thermo Fisher Scientific), as per the protocol supplied by the manufacturer. The assay mixtures for reverse transcriptase reactions contained 1 µg RNA, 0.112 µM oligo(dT), 0.5 mM dNTP, 200 units of High Capacity RT (Applied Bio-Systems) in 1× RT buffer. cDNA was stored at –20°C.

### 2.6. Quantitative real-time polymerase chain reaction (RT-qPCR) analyses

The expression patterns of clock genes, osteo/chondrogenic transcription factors and genes coding for cartilage ECM structural proteins were determined using RT-qPCR by relative quantification using the 2^−ΔΔCt^ method. Primer pairs were obtained from Integrated DNA Technologies (Coralville, IA, USA). For sequences of custom-designed primer pairs please see *Table S1* in the Supporting Information. SYBR Green-based RT-qPCR reactions were set up using the GoTaq qPCR Master Mix (Promega) and 20 ng input cDNA per each 10-μL reaction. Reactions were run in a QuantStudio 3 Real-Time PCR System (Thermo Fisher Scientific) using the following standard thermal profile: activation and initial denaturation at 95°C for 2 minutes, followed by 40 cycles of denaturation at 95°C for 3 sec, annealing and extension at 60°C for 30 sec, and then final extension at 72°C for 20 sec. Data was collected during the extension step. Amplification was followed by a melt curve stage consisting of 3 steps: denaturation at 95°C for 15 sec, annealing at 55°C for 1 min, followed by a dissociation step at 0.15°C/sec increments between 55°C and 95°C. Data collection was enabled at each increment of the dissociation step. Amplification data were analysed using the QuantStudio Design and Analysis Software (version 1.5.1) and exported data were processed using Microsoft Excel (version 2108).

Based on previous literature, three reference genes were analysed for stability during *in vitro* chondrogenesis at each time point as follows: peptidylprolyl isomerase A (*PPIA*), 60S ribosomal protein L13 (*RPL13*) and tyrosine 3-monooxygenase/tryptophan 5-monooxygenase activation protein zeta (*YWHAZ*). Microsoft Excel was employed to determine the optimal normalising gene between the chosen reference genes based on their expression stability values. Real-time qPCR data for each gene of interest were normalised to the most stable reference gene levels in the same sample.

### 2.7. Gene expression data analysis and cosine fits

Presence of circadian rhythmicity was determined by an extra-sum-of-squares F test (α = 0.05) to test the hypothesis that gene expression was best described by a cosine curve + sloping line, rather than the null hypothesis of a sloping line alone. Curve fitting and F testing was conducted in MATLAB (v 2019a, MathWorks, Natick, MA, USA). A sloping line and single cosine were fitted to all replicate values in a single fit, and returned an intercept and slope for the line, and a period (bounds set to 20–28 hours), amplitude, acrophase and mesor if a significant cosine fit was found (see *Table 2*). Data is plotted as mean ± SD, with non-synchronised controls (grey) and cells subjected to mechanical loading (blue), including the linear and cosine fits.

### 2.8. Validation of cartilage ECM production in micromass cultures by metachromatic staining

Micromass cultures were set up onto the surface of 30-mm round glass cover slips (Menzel-Gläser, Menzel GmbH, Braunschweig, Germany) placed into 6-well culture plates. For qualitative and semi-quantitative evaluation of cartilage matrix production, cultures were exposed to the mechanical loading scheme on each day from day 1 until day 6 for 1 hour, and were stained on day 6 after the last 1-hour stimulus with dimethyl-methylene blue (DMMB; pH 1.8; Merck) metachromatic dyes as previously described ^15^. Some colonies received 5 µM of LDS for the duration of the mechanical load on each culturing day. The optical density values of DMMB-stained specimens were determined from cultures in 3 independent, biological replicate experiments using a MATLAB image analysis application. Cartilage nodules rich in metachromatic cartilage ECM were defined by an approximate range of values in the RGB colour space and the pixels were counted.

### 2.9. Simple Western Wes immunoassay for protein expression

Micromass cultures (either following mechanical stimuli or untreated controls) collected every 8 hours from 24 to 72 hours post-stimulation on day 6 were lysed in radioimmunoprecipitation (RIPA) lysis buffer (Pierce, Thermo Fisher Scientific; Cat. No.: 89900) containing protease inhibitors (Sigma-Aldrich; Cat. No.: P8340) using an ultrasonic processor (Cole-Parmer, Vernon Hills, IL, USA). Equal amounts of protein (3 µg) were loaded into 12–230 kDa separation modules (Protein Simple, Bio-Techne, Minneapolis, MN, USA; Cat. No.: SM-W004) and analysed using the Protein Simple Wes System (Protein Simple, Bio-Techne; Cat. No.: 004–600) with the Anti-Rabbit Detection Module (Protein Simple, Bio-Techne; Cat. No.: DM-001), following the manufacturer’s instructions. Briefly, samples were diluted to an appropriate concentration (1 μg/μL) in sample buffer (‘10× Sample Buffer’ from the Separation Module), then mixed with Fluorescent Master Mix 1:4 and heated at 95 °C for 5 min. The samples, the blocking reagent (antibody diluent), the primary antibodies, the HRP-conjugated secondary antibodies, and the chemiluminescent substrate were added to the plate. The default settings of the device were as follows: stacking and separation at 395 V for 30 min; blocking reagent for 5 min; primary and secondary antibodies both for 30 min; luminol/peroxide chemiluminescence detection for 15 min (exposure times were selected for the antibodies between 1 and 512 s). The electropherograms were visually checked, and the automatic peak detection was manually corrected if required. Wes data are obtained as virtual blots in which the molecular weight and signal intensity are presented. Results in the form of electropherograms are also obtained with this approach. Molecular weights associated with the bands are presented, and the area under the curve in these plots correspond to the total chemiluminescence intensities. The analysis was carried out on three replicates (*N* = 3). *Table 1* depicts the primary antibodies and their dilutions (all at 1:25) used in the Wes system.

**Table 1.** Antibodies and their dilutions used in the Wes system

| Antibody | Cat. No. | Vendor | Host | Dilution |
| --- | --- | --- | --- | --- |
| Aggrecan | ABT545 | Merck Millipore | Rabbit polyclonal | 1:25 |
| BMAL1 | ab93806 | Abcam | Rabbit polyclonal | 1:25 |
| CRY1 | ab104736 | Abcam | Rabbit polyclonal | 1:25 |
| PER3 | ab177482 | Abcam | Rabbit polyclonal | 1:25 |
| RUNX2 | 8486S | Cell Signaling Technology | Rabbit monoclonal | 1:25 |
| SOX9 | ab3697 | Abcam | Rabbit polyclonal | 1:25 |
| GAPDH | ab9485 | Abcam | Rabbit polyclonal | 1:25 |

**Table 2.** Cosinor analysis parameters of core clock and chondrogenic marker gene expression in chondrogenic cultures following mechanical load (**A**) and in non-stimulated control cultures (**B**). *P*, probability value; R^2^, coefficient of determination

|  | *P* | Mesor |  | Amplitude | Acrophase | Period | *R*<sup>2</sup> |
| --- | --- | --- | --- | --- | --- | --- | --- |
|  |  | Intercept | Slope |  |  |  |  |
| *BMAL1* | <0.001 | 1.500 | -0.016 | -0.121 | 22.461 | 21.918 | 0.951 |
| *REV-ERB* | <0.05 | 1.330 | 0.006 | -0.600 | 20.456 | 20.000 | 0.461 |
| *PER2* | <0.001 | 1.360 | -0.013 | -0.178 | 20.114 | 22.046 | 0.897 |
| *PER3* | <0.01 | 1.145 | -0.007 | 0.240 | 28.000 | 25.007 | 0.731 |
| *CRY1* | <0.05 | 1.402 | -0.012 | -0.152 | 25.743 | 20.000 | 0.809 |
| *CRY2* | 0.32 | 1.278 | -0.010 | 0.086 | 16.331 | 20.000 | 0.610 |
| *SOX6* | <0.01 | 2.757 | -0.037 | 0.556 | 12.211 | 20.000 | 0.792 |
| *SOX9* | <0.01 | 3.306 | -0.038 | 1.096 | 10.163 | 22.238 | 0.670 |
| *ACAN* | <0.01 | 2.030 | -0.017 | 0.380 | 11.668 | 21.596 | 0.758 |
| *COL2A1* | 0.09 | 1.165 | -0.003 | -0.313 | 18.668 | 20.000 | 0.373 |
| *RUNX2* | 0.15 | 2.093 | -0.023 | 0.243 | 10.450 | 28.000 | 0.664 |

**B. Non-stimulated (control)**
|  | *P* | Mesor |  | Amplitude | Acrophase | Period | *R*<sup>2</sup> |
| --- | --- | --- | --- | --- | --- | --- | --- |
|  |  | Intercept | Slope |  |  |  |  |
| *BMAL1* | 0.89 | 1.069 | 0.000 | 0.053 | 8.595 | 28.000 | 0.065 |
| *REV-ERB* | 0.61 | 0.980 | 0.001 | -0.038 | 9.172 | 28.000 | 0.189 |
| *PER2* | 0.91 | 0.944 | 0.001 | 0.026 | 20.138 | 20.000 | 0.065 |
| *PER3* | 0.98 | 1.040 | 0.000 | 0.019 | 16.282 | 21.919 | 0.020 |
| *CRY1* | 0.55 | 1.012 | 0.001 | 0.064 | 15.970 | 21.246 | 0.250 |
| *CRY2* | 0.92 | 1.006 | -0.001 | -0.029 | 17.095 | 20.997 | 0.074 |
| *SOX6* | 0.89 | 1.536 | 0.023 | 0.402 | 11.812 | 28.000 | 0.574 |
| *SOX9* | 0.72 | 0.930 | 0.000 | -0.039 | 10.010 | 28.000 | 0.141 |
| *ACAN* | 0.79 | 0.911 | 0.002 | 0.034 | 22.502 | 20.000 | 0.201 |
| *COL2A1* | 0.92 | 1.005 | 0.001 | 0.043 | 17.526 | 21.491 | 0.053 |
| *RUNX2* | 0.77 | 0.862 | 0.003 | -0.033 | 11.199 | 23.373 | 0.360 |

### 2.10. Immunohistochemistry for collagen type II expression

Immunohistochemistry was performed on micromass cultures seeded onto the surface of glass coverslips to visualise collagen type II expression. After the loading scheme on day 6, cultures were fixed in 4% paraformaldehyde (Sigma-Aldrich, Merck) for 30 min and washed in distilled water. Nonspecific binding sites were blocked with PBS supplemented with 3% BSA (Amresco Inc., Solon, OH, USA) and 10% goat serum (Vector Labs, Inc., Burlingame, CA, USA) for 30 min, at room temperature. Thereafter, samples were incubated with anti-collagen type II polyclonal antibody produced in rabbit (Novus Biologicals, Bio-Techne, Minneapolis, MN, USA; Cat. no: NB600-844) at a dilution of 1:250 in PBS supplemented with 1% BSA and 3% goat serum, at 4°C overnight. On the following day, after washing three times with PBS, primary antibodies were visualized with diamino benzidine (DAB) chromogen reaction (Sigma-Aldrich, Merck), and samples background-stained with haematoxylin (Amresco). For the DAB chromogen reaction, biotinylated anti-rabbit antibody (Vector Labs; Cat. no: BA-1000-1.5) was used on the samples for 1 hour at a dilution of 1:1000 in PBS, followed by additional wash steps. Then, Extravidin (Sigma-Aldrich, Merck) was applied on the samples for another 1 hour at a dilution of 1:1000 in PBS. After further washing, the DAB reaction was performed, and samples were stained with haematoxylin to visualize cell nuclei. Finally, slides were mounted with DPX medium and coverslipped (Sigma-Aldrich). Photomicrographs of the micromass cultures were taken using an Olympus BX53 camera on a Nikon Eclipse E800 microscope (Nikon Corporation, Tokyo, Japan). Reactions were carried out on three biological replicates (*N* = 3), and photomicrographs of five different visual fields were examined for each sample.

### 2.11. Statistical analyses

All experiments were performed 3 times (*N* = 3 biological replicates). For RT-qPCR and protein assays, one representative data set is shown out of 3 parallel experiments, each displaying identical dynamics. Microsoft Excel was employed to perform group analysis. For the assessment of cartilage ECM production, optical density values are shown as means ± standard error of the mean (SEM); statistical differences were determined using paired Student’s *t*-test. For RT-qPCR data analysis, expression data of each time point were compared to the previous data point and statistical differences were determined using paired Student’s *t*-test. As indicated above, presence of circadian rhythmicity was determined by an extra-sum-of-squares F test comparing a sloping line against a cosine + line model (α = 0.05).

## 3. Results

### 3.1. Intermittent cyclic mechanical load augmented metachromatic cartilage matrix production in micromass cultures in a circadian clock-dependent manner

We first checked whether uniaxial cyclic compressive force administered to micromass cultures every day between days 1 and 6 for 60 min at the same time of each day influenced metachromatic cartilage ECM production by day 6. Following DMMB staining, image analysis confirmed that mechanical stimulation (MS) massively enhanced matrix production to 233% (±3.96%; *P* < 0.001) compared to the control (*Fig. 2A*), without influencing cell viability (*Fig. S1* in the Supporting Information). Another set of cultures were exposed to 5 µM of LDS for 1 hour every day; perturbing the circadian clock by this treatment did not interfere with cartilage ECM production by day 6 (91.19 ± 11.4%; *P* = 0.26). Combined exposure to MS and LDS for 60 min for 6 days resulted in a significant (155 ± 1.89%) increase in metachromatic matrix production (*P* < 0.001 *vs*. the control), but it reduced ECM deposition compared to MS alone (*P* < 0.001).

**Figure 2.**
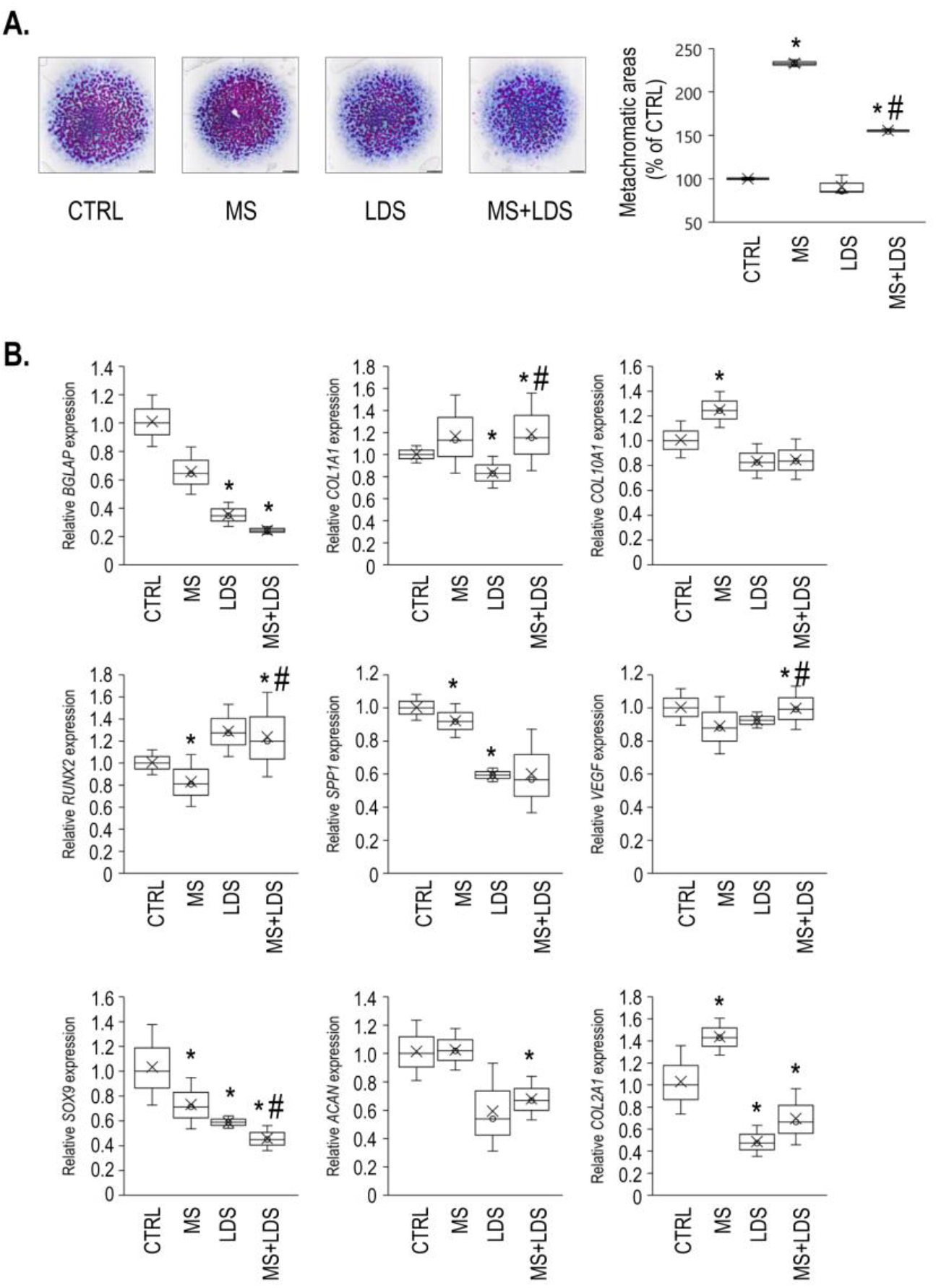
Mechanical load administered to micromass cultures for 60 min on days 1 through 6 increased chondrogenesis and metachromatic cartilage ECM production by day 6 in a circadian clock-dependent mechanism. **A**. Metachromatic cartilage ECM accumulation during chondrogenesis in control *vs*. mechanical load-induced cultures, with or without longdaysin (LDS) treatment, as determined by DMMB (qualitative) staining assay. Original magnification was 2×. Scale bar, 1 mm. The bar chart shows the results of a MATLAB-based image analysis of metachromatic areas in mechanically loaded *vs*. control cultures, with or without LDS treatment. Data are expressed as mean ± SEM. Statistical significance (*P* < 0.05) between mechanically stimulated and control cultures is indicated by asterisks (*), and between LDS-treated and untreated samples by a hash sign (#). Representative data out of 3 independent experiments. **B**. Cartilage-specific, osteogenic, and chondrocyte hypertrophy marker gene expression on day 6 following the mechanical loading scheme, with or without treatment with the circadian clock regulator LDS. Data are expressed as the mean ± SD relative to the control and normalised against the reference gene *RPL13*. Statistical significance (*P* < 0.05) between gene expression levels is indicated by asterisks (relative to vehicle control) or a hash sign (between LDS-treated and untreated cultures). CTRL, control; MS, mechanical stimuli; LDS, longdaysin

To assess whether the observed changes in cartilage ECM production were detectable at the molecular level, the gene expression patterns of key osteo-chondrogenic marker genes, including the master transcription factor *SOX9*, the alpha-1 chain of collagen type II (*COL2A1*), and aggrecan (*ACAN*) were tested using RT-qPCR (*Fig. 2B*). *COL2A1* mRNA expression was significantly upregulated (1.4-fold ± 0.093; *P* = 0.009) in cultures exposed to rhythmic MS for 6 consecutive days, relative to the control. *ACAN* levels remained unchanged, whereas *SOX9* expression was downregulated following MS in 6-day-old colonies compared to the control. When the circadian clock modulator compound LDS was applied for 1 hour on every culturing day for 6 days, all three osteo-chondrogenic markers displayed a significant downregulation of their expression. Combined treatment with LDS and cyclic mechanical load resulted in lower gene expression levels for the studied chondrogenic marker genes relative to the untreated control (*Fig. 2B*). The loading scheme enhanced collagen type II protein expression in micromass cultures compared to the unstimulated control cultures, as revealed by collagen type II immunohistochemistry (*Fig. 3*).

**Figure 3.**
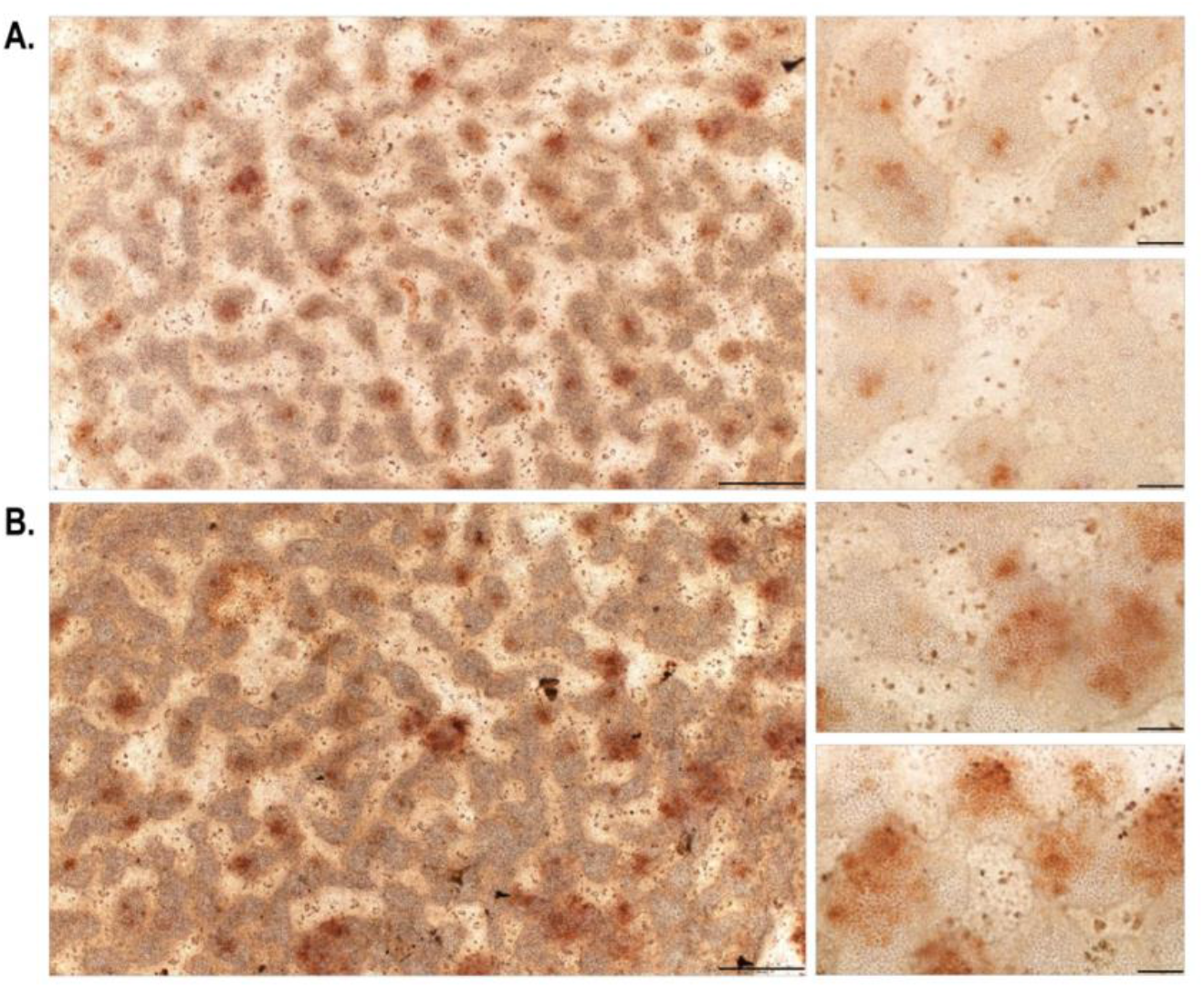
Collagen type II immunohistochemistry visualised by DAB chromogen reaction in 6-day-old unstimulated chondrifying micromass cultures (**A**) compared to colonies that received the mechanical loading regime for 6 consecutive days (60 minutes daily) (**B**). Scale bars, 500 μm (main images on the left) or 100 μm (smaller inserts on the right). The brown deposit represents immunopositive signals for collagen type II within the chondrogenic nodules of the micromass cultures. Representative photomicrographs are shown.

We also looked at the gene expression profiles of osteogenic and chondrocyte hypertrophy markers, namely *COL1A2* (type I collagen, alpha chain), *COL10A1* (type X collagen, alpha chain), *BGLAP* (bone gamma-carboxyglutamate protein, osteocalcin), *VEGF* (vascular endothelial growth factor), *SPP1* (secreted phosphoprotein, osteopontin), and *RUNX2* (RUNX family transcription factor 2) (*Fig. 2B*). Except for *COL10A1*, a marker of hypertrophy (a sign of maturation in this model), which was significantly upregulated following the loading regime (1.2-fold ± 0.057; *P* = 0.037), the osteogenic markers (*BGLAP, RUNX2, SPP1*, and *VEGF*) were downregulated following 6-day-long mechanical stimuli *versus* the control. LDS treatment (with or without mechanical stimulation) downregulated the osteogenic markers except *RUNX2* (*Fig. 2B*).

Taken together, these results suggest that rhythmic mechanical stimuli may have enhanced chondrogenesis and ECM deposition, at least partially, *via* a circadian clock-dependent mechanism.

### 3.2. The molecular clock genes exhibited a synchronised oscillatory pattern during early chondrogenesis following mechanical load

Transcripts for the clock genes *BMAL1, PER2/3, CRY1/2*, and *REV-ERB*, known to control the molecular clockwork, were previously detected in chondrifying cells of micromass cultures, indicating that these genes are already expressed and functional during early chondrogenesis ^15^. To establish whether the clock genes exhibited a rhythmic expression over time following the loading scheme, micromass cultures were exposed to cyclic intermittent uniaxial cyclic mechanical load for 1 hour at the same time of each culturing day between days 1 and 6. Samples for total RNA isolation were collected at 24 hour after the last loading regime (time point 0) and then at every 8 hours for the period of 24–72 hours to identify oscillatory patterns in clock gene expression. Time courses of the relative quantity (RQ) and Relative Standard Deviation (RSD) values for the core clock genes (after being normalised to *RPL13*, the most stable reference gene tested) are shown in *Table S2* in the Supporting Information. As indicated, the normalised relative quantity values for each gene of interest at the 24-hour time point was set to 1.0.

We detected a synchronised oscillatory expression pattern for the majority of the clock genes studied over the investigated 72-hour period as determined by a significant fit to a non-linear cosinor model (*Fig. 4*). There was an approx. 6-hour phase angle between the positive phase and negative phase of the TTFL (i.e., *BMAL1 vs. CRY1* and *PER3*) with periods between 20–24 hours. The parameters of the observed circadian expression patterns for 6-day-old chondrogenic cultures synchronised with daily mechanical load are shown in *Table 2*.

**Figure 4.**
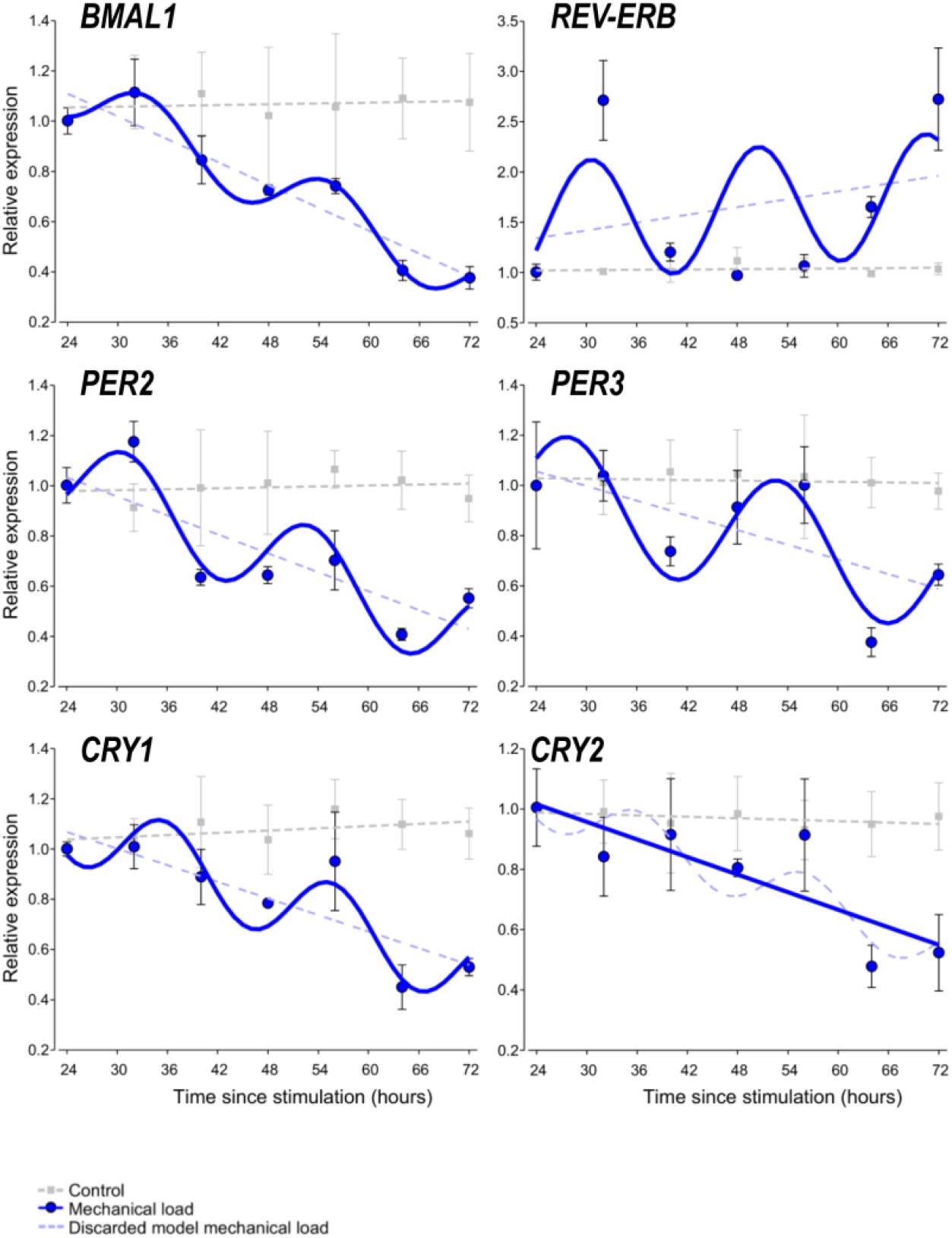
Circadian rhythm dynamics in clock gene expression in 6-day-old differentiating embryonic limb bud-derived chondroprogenitor cells of micromass cultures following synchronisation with uniaxial cyclic mechanical load. Quantitative RT-PCR analyses followed by cosine fits showing temporal expression profiles of the clock genes collected every 8 hours between 24–72 hours post-synchronisation with mechanical load (solid blue line or curve) *versus* non-stimulated control cultures (broken grey lines) collected at the same time points. Each plot also includes the discarded model (i.e., a broken straight blue line if the cosine fit was significant, or a broken blue sinusoidal curve if the cosine fit was not significant). Data are expressed as the mean of transcript levels ± SD relative to the 24-hour time point and normalised against the reference gene *RPL13*. Representative data are shown out of 3 independent experiments, each exhibiting similar patterns of gene expression profiles

### 3.3. The chondrogenic marker genes also show a circadian expression pattern during chondrogenesis

Chondrogenic differentiation in embryonic limb bud-derived micromass cultures is controlled by the osteo-chondrogenic transcription factors *SOX6, SOX9*, and *RUNX2*. These transcription factors are upstream regulators of genes encoding cartilage ECM components such as *ACAN* and *COL2A1*. In order to establish whether the expression of these marker genes also showed an oscillatory pattern, their transcript levels were analysed by RT-qPCR, and then the expression values were fitted with the cosine function to reveal circadian rhythms in their expression (*Fig. 5*). The cosinor parameters of the observed circadian expression patterns for the marker genes in chondrogenic cultures are summarised in *Table 2*.

**Figure 5.**
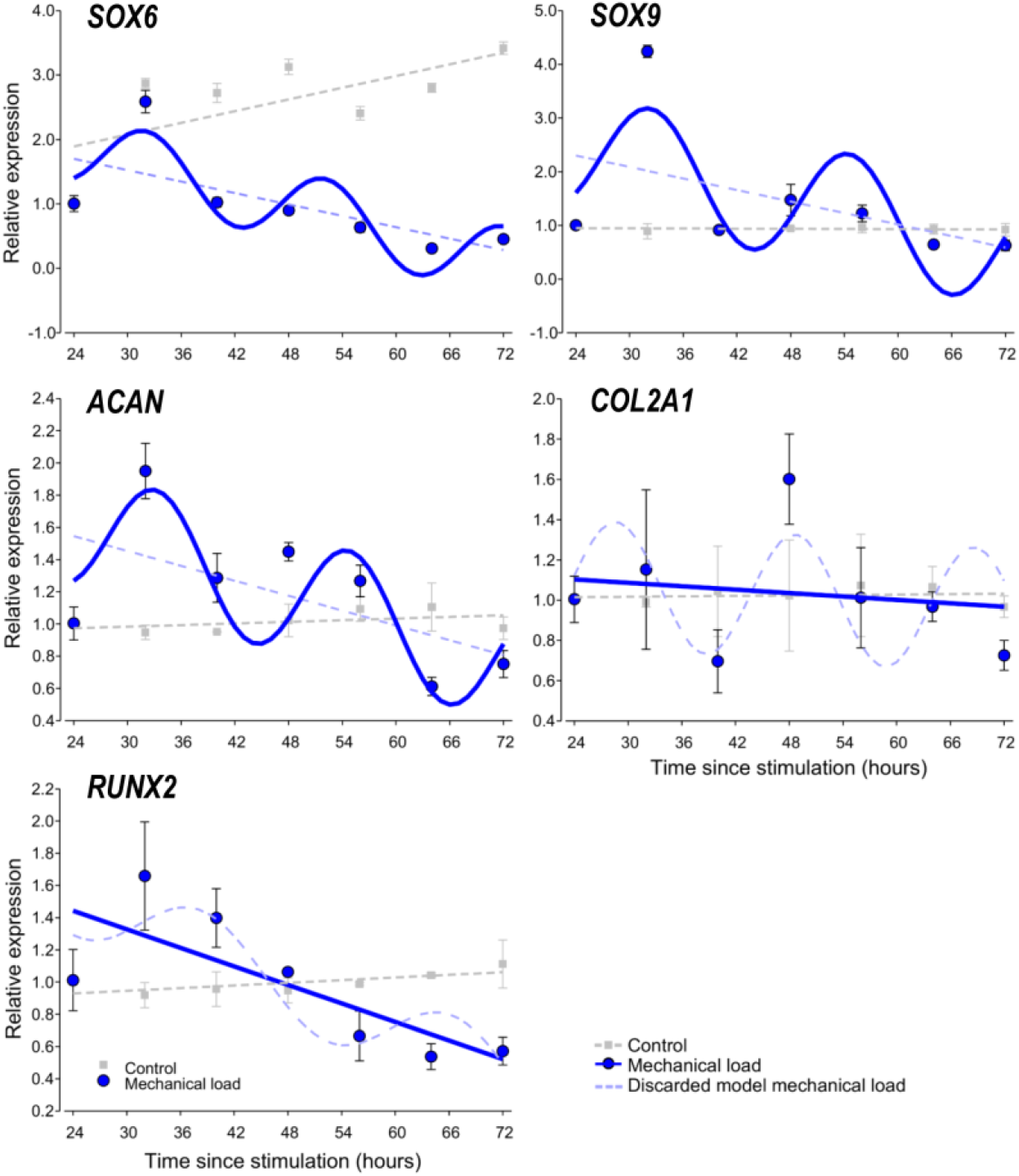
Circadian rhythm dynamics in clock gene expression in 6-day-old differentiating embryonic limb bud-derived chondroprogenitor cells of micromass cultures following synchronisation with uniaxial cyclic mechanical load. Quantitative RT-PCR analyses followed by cosine fits showing temporal expression profiles of chondrogenic marker genes collected every 8 hours between 24–72 hours post-synchronisation with mechanical load (solid blue line or curve) *versus* non-stimulated control cultures (broken grey line) collected at the same time points. Each plot also includes the discarded model (i.e., a broken straight blue line if the cosine fit was significant, or a broken blue sinusoidal curve if the cosine fit was not significant). Data are expressed as the mean of transcript levels ± SD relative to the 24-hour time point and normalised against the reference gene *RPL13*. Representative data are shown out of 3 independent experiments, each exhibiting similar patterns of gene expression profiles

In 6-day-old chondrogenic micromass cultures synchronised by the cyclic uniaxial mechanical loading scheme, *SOX6, SOX9*, and *ACAN* displayed oscillatory expression patterns (*Fig. 5*). In contrast, no circadian pattern could be detected for *RUNX2* and *COL2A1*. Time courses of the relative quantity and relative SD values for the core clock genes (after being normalised to *RPL13*, the most stable reference gene tested) are shown in *Table S2* in the Supporting Information. These findings suggest a circadian regulation of chondrogenic marker genes in micromass cultures during chondrogenesis as a result of intermittent mechanical load, and that the two key transcription factors *SOX9* and *RUNX2* may be differentially regulated by the molecular clock following daily entrainment with mechanical load.

### 3.4. Core clock components and chondrogenic markers display rhythmic expression patterns at the protein level

Having established that most of the core clock genes and some of the chondrogenic marker genes analysed displayed a robust circadian expression pattern at the transcript level following the mechanical load *versus* unstimulated control cultures, we then looked at whether the above patterns were reflected at the protein level. For protein detection in the micromass cultures, we employed a capillary-based Simple Western Wes immunoassay platform which offers key advantages including automation, better reproducibility, and more reliable quantification ^29^. We monitored protein expression patterns for the following core clock proteins at the same time points as for the transcript analysis: BMAL1, CRY1, and PER3. All three clock proteins displayed a more robust circadian rhythmic expression (∼24 hours between two peaks) following the loading scheme compared to the control (*Fig. 6*). These data indicate that the intermittent mechanical loading scheme has induced a rhythmic expression pattern of the clock proteins in chondrogenic cultures. Furthermore, the protein levels of CRY1, which was relatively weakly expressed in unstimulated control cultures, were higher at all time points in cultures that received mechanical loading.

**Figure 6.**
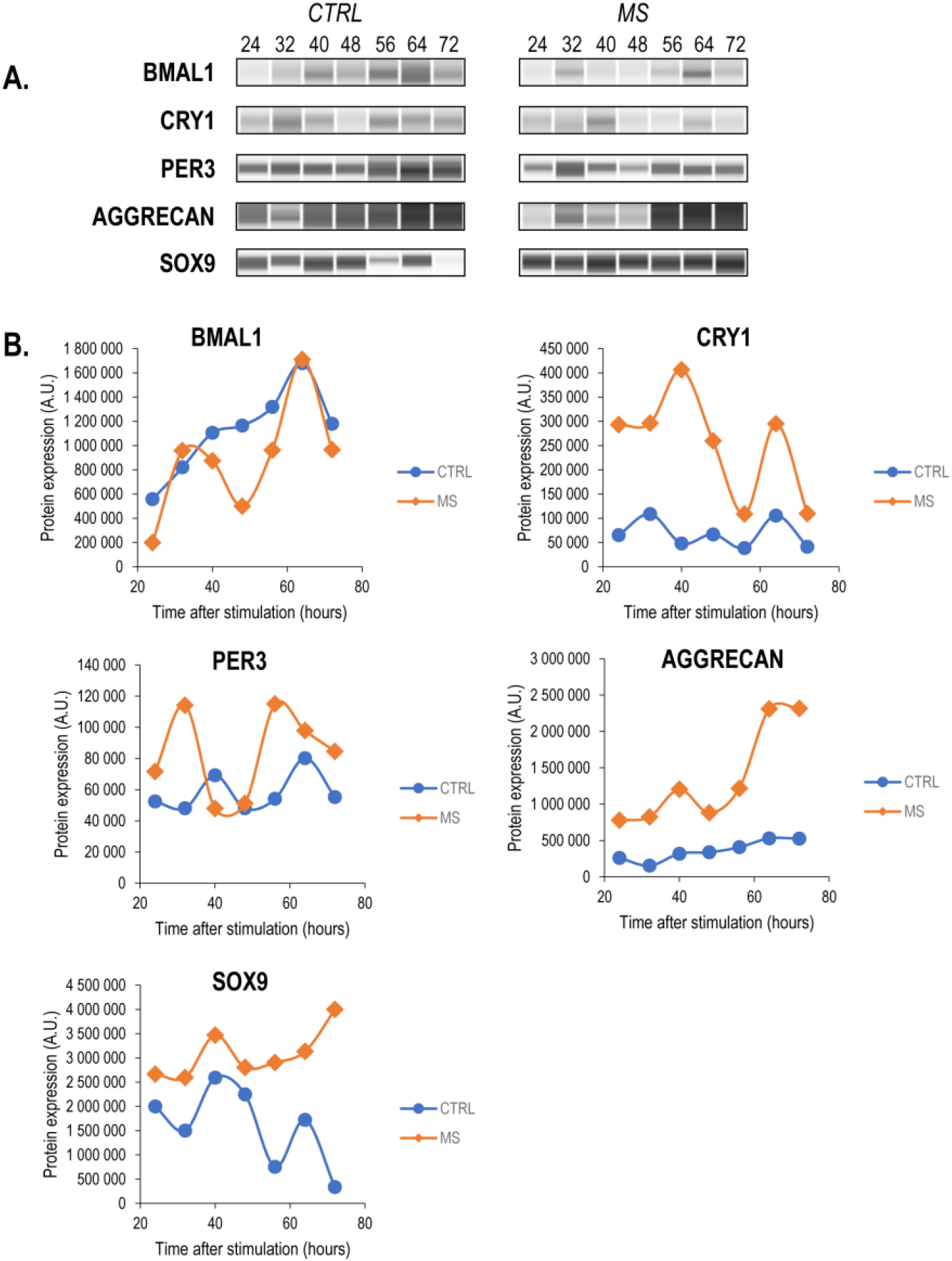
Wes immunoassay data (quantitative and software-generated capillary lane images) for the core clock proteins BMAL1, CRY1, and PER3, the cartilage ECM structural component aggrecan core protein, and the master chondrogenic transcription factor SOX9, in mechanically stimulated (MS) micromass cultures *versus* the control (CTRL) at 24–72 hours post stimulation. (**A**) Representative Wes immunoassay electropherograms are shown displaying the main trends observed on the individual biological replicates. (**B**) Quantitative analysis of the Wes immunoassay results. Uncropped electropherograms are shown in *Fig. S2* in the Supplementary material

We also looked at the protein level expression pattern of those two chondrogenic markers (aggrecan and SOX9) whose transcripts displayed a sinusoidal pattern. In these cases, in addition to having observed a rhythmic protein expression pattern (∼24-hour period between two peaks) following mechanical stimuli, both aggrecan and SOX9 were more abundantly expressed compared to untreated control cultures (*Fig. 6*). We also aimed to analyse the protein expression pattern of the osteogenic transcription factor RUNX2, but we could not detect specific signals at the expected molecular weight (*Fig. S2*). The uncropped Wes electropherograms for each run shown in *Fig. 6* are shown in *Fig. S2* in the Supporting material.

## 4. Discussion

This is the first report to show that cyclic uniaxial mechanical stimulation augments chondrogenic differentiation in primary embryonic limb bud-derived micromass cultures by promoting rhythmic expression of the circadian molecular clock. When biochemical perturbation of the clock was combined with mechanical stimulation, a reduction was detected in the mechanical stimulation-driven enhanced chondrogenesis. These observations show that mechanical stimulation can act as peripheral *Zeitgebers* of circadian clock during early foetal development.

Timing of peripheral circadian clocks, which are expressed in almost all nucleated cells of multicellular organisms, is co-ordinated by the central pacemaker clock in the central nervous system. These peripheral clocks in turn control rhythms in local molecular and cellular processes. Transcriptional analyses have shown that a large proportion of the genome is clock-controlled; depending on the experimental setting, up to 20% of protein-encoding genes display circadian oscillatory patterns ^30,31^. In addition to transcriptional changes, circadian regulation of cell physiology arises from the rhythmic control of post-transcriptional processes, including RNA splicing, protein translation, and post-translational modifications. Indeed, a recent study confirmed that 145 of the 1177 identified proteins (12%) displayed 24-hour rhythmic changes in the proteome of femoral head cartilage ^32^.

Although the master clock is entrained by light, and peripheral clocks are normally entrained by the central pacemaker, there are other cell and tissue-specific factors (*Zeitgebers*) that can synchronise these peripheral clocks. The internal factors that are known to be able to act as timing cues for peripheral clocks include body temperature and the sympathetic nervous system, circulating factors such as glucocorticoids, cytokines, melatonin, and molecular nutrients sensors for e.g., glucose, temperature, redox status and AMPK that impact on clock gene expression ^33-35^. Exposing cultured cells, including chondrogenic progenitor cells, to a pulse of serum *in vitro* synchronizes robust 24-hour transcription cycles ^15,30^. This is thought to be driven by factors in the serum, including glucocorticoids, which act as physiological timing cues that synchronise the individual cellular clocks in the culture to the same phase leading to a culture-wide, synchronised circadian rhythm ^36,37^. We here show that mechanical stimulation also acts as a timing cue that synchronises clock gene expression in chondrogenic progenitor cells *in vitro*, that enhances chondrogenesis. The physiological relevance of this for the human circadian timing system is that mechanical movement of cartilage may synchronise local peripheral clocks in this tissue. When this mechanical movement is mistimed (e.g., when being active at night during shift work), this non-photic timing cue that can be ‘uncoupled’ from light entrained timing cues leading to circadian desynchrony between central and peripheral timing.

Circadian desynchrony is most frequently studied in the context of metabolic timing cues that desynchronise peripheral clocks from the central, light-entrained clock. Feeding can independently synchronize peripheral tissue clocks in the liver and kidneys; food consumed at the wrong time of day, resulting in changes in the circadian timing of metabolic cues such as glucose levels, redox status and AMPK that act as timing cues for peripheral clocks, may therefore lead to misalignment in clock timing ^38^. Irregular sleep and eating schedules can misalign clocks in metabolic organs, leading to obesity and diabetes ^30^. Understanding the molecular actions of mechanical stimulation on the circadian clock, and the cell and tissue-specific *Zeitgebers* may provide insight into the mechanism by which this novel peripheral timing cue acts on circadian clocks that are involved in tissue homeostasis in developing and mature tissues.

Given its location and poor circulation due to its avascular nature, the molecular clock in chondrocytes of articular cartilage is unlikely to be directly entrained by light and metabolic cues. In this work, we have shown, for the first time, that properly adjusted mechanical stimuli that chondrogenic cells are also exposed to *in vivo* are able to entrain the molecular clockwork during chondrogenic differentiation. The impact of mechanical signalling on developing and mature cartilage is well appreciated, albeit the specific mechanotransduction pathways are not completely understood. We have previously shown that mechanical stimuli augment *in vitro* cartilage formation *via* promoting both differentiation and matrix production of chondrogenic cells, through the opposing regulation of the PKA/CREB-Sox9 and the PP2A signalling pathways in the embryonic limb bud-derived micromass model ^14^. Mechanosensitive ion channels such as transient receptor potential vanilloid 4 (TRPV4), PIEZO1, and PIEZO2 play important functional roles in chondrocyte mechanosensation ^39^. While several ion channels are expressed in chondrocytes that regulate calcium signalling ^40^, TRPV4 and PIEZOs are two of the most relevant ion channels in chondrocytes in this context, both have been shown to improve cartilage and bone formation, and by mediating the anabolic effects of mechanical loading ^41^. However, extracellular calcium and membrane calcium channels were not found to be involved in hyperosmolarity-induced clock resetting in mouse cartilage ^42^, and the mTORC2-AKT-GSK3β pathway is hypothesised to act as a convergent mechanism mediating clock entrainment elicited by mechanical loading and hyperosmolarity ^42^. Nevertheless, specific pathways activated upon different external stimuli may be dependent on a number of factors such as the exact nature of the stimuli and the differentiation phase of the cell.

Mechanical cues are also pivotal in modulating the chondrogenic differentiation of MSCs. Dynamic compressive loading has been previously shown to stimulate MSC chondrogenesis, with reports describing upregulated SOX9 expression, and increased glycosaminoglycan and collagen type II production ^6^.

Since certain components (e.g., *CLOCK*) of the molecular clockwork were shown to be mechanosensitive ^19^, we attempted to synchronise the clock in differentiating chondrocytes using a cyclic mechanical loading scheme. Rhythmic mechanical stimulation was shown to have the ability to entrain human MSCs, which represents a novel clock synchronization approach independent of chemical or temperature cues ^22^. Indeed, uniaxial cyclic compressive force (approx. 0.6 kPa, 0.05 Hz) administered to cells of chondrifying micromass cultures on every culturing day for 6 days (for 60 min each day) synchronised the expression of the core clock genes and cartilage marker genes, and resulted in augmented cartilage ECM deposition, as revealed by metachromatic staining with DMMB, as well as collagen type II immunohistochemistry. We also observed that differentiating chondrocytes responded differently to circadian synchronising signals; in serum shock-synchronised cells, *BMAL1, CRY1*, and *PER2* displayed sinusoidal expression patterns ^15^, whereas mechanical stimuli caused *BMAL1, CRY1, PER2, PER3*, and *REV-ERB* to follow a circadian pattern. Additionally, whilst *COL2A1* showed a sinusoidal expression pattern in serum-shocked cultures ^15^, it did not become rhythmic following entrainment by MS.

We also analysed the protein expression of the core clock components and established that the intermittent mechanical loading scheme triggered BMAL1, CRY1, and PER3 proteins to display rhythmic (∼24-hour period) changes in their abundance in chondrogenic micromass cultures. In a previous study which investigated daily changes of protein abundance in mouse femoral head articular cartilage by performing a 48-hour time-series LC-MS/MS analysis, 145 proteins (out of the 1,177 proteins identified) were shown to display rhythmic changes ^32^; however, none of these proteins were observed in that dataset. The rhythmic expression of BMAL1 and CRY1 proteins has previously been reported in ATDC5 cells differentiated into chondrocytes; however, in that study, the expression was monitored for 48 hours only ^43^. To the best of our knowledge, this is the first report on the circadian modulation of the core clock proteins BMAL1, CRY1, and PER3 in chondrifying micromass cultures as a result of cyclic intermittent mechanical loading.

It appears that chondrogenic cells cultured in a mechanical environment exhibit a different profile of the molecular circadian clock compared to synchronisation by glucocorticoids and growth factors present in high levels of foetal serum. With regards to the negative loop of the core clock, *CRY2* also displayed a sinusoidal pattern but it could not be fitted with a cosine curve (see *Fig. 4*, and *Table 2*). The fact that *PER2* also showed oscillatory expression following MS is in line with the pattern identified in human bone marrow-derived MSCs (BMSC) and dental pulp stem cells (DPSC), where *PER2* was also found to be responding to rhythmic mechanical stretch ^22^. In a similar way, the rhythmic expression of *REV-ERB* elicited by MS in our model also coincides with the sinusoid pattern in bone marrow-derived MSC and dental pulp stem cells exposed to mechanical stress ^22^.

We observed that the mechanical loading regime significantly increased cartilage ECM production and upregulated *COL2A1* expression both at the mRNA and the protein level. At the same time, *SOX9* expression was downregulated, probably indicating that the dynamic mechanical microenvironment accelerated the pace of chondrogenic differentiation; the observed lower transcript levels of *SOX9* are characteristic to more mature chondrocytes ^44^. In contrast, we found that SOX9 and aggrecan protein levels were higher following the mechanical loading scheme compared to unstimulated controls, which explains the observed chondro-stimulatory effects. Furthermore, a more rhythmic protein expression pattern was observed in case of SOX9 and aggrecan in cultures exposed to mechanical stimuli.

The observation that the cultures exposed to the loading scheme favoured chondrogenic differentiation is also substantiated by the fact that most osteogenic marker genes examined (i.e., *BGLAP, RUNX2, SPP1, VEGF*) displayed attenuated expression levels following the loading scheme. RUNX2 protein was expressed at very low levels only (under the detection threshold of Simple Western Wes), further supporting blocked osteogenic pathways in micromass cultures. To check whether interference with the normal circadian rhythm generated by the loading regime in chondrogenic cultures had any observable effects on the parameters measured, we used the clock modulator compound longdaysin. When LDS was applied in addition to mechanical load, a moderately enhanced metachromatic ECM production was observed, despite reduced chondrogenic marker gene expression. This could have been caused by cross-talk with other, circadian clock-independent signalling pathways (i.e., PKA/CREB-SOX9 and PP2A ^14^) through the regulation of glycosaminoglycan deposition at the post-transcriptional level.

In terms of protein expression analysis, we observed that GAPDH levels also followed a rhythmic expression pattern in micromass cultures receiving the loading scheme *versus* the control (see *Fig. S2*). GAPDH was also identified in the mouse femoral head articular cartilage 48-hour time-series LC-MS/MS analysis ^32^, but it was not classified as a rhythmically expressed protein due to high *P* values (*P* > 0.05). Nevertheless, our finding may be relevant when choosing normalising genes/proteins for data analysis in a particular setting.

The observed differences and similarities between our model and the MSC-based models described above can be attributed to (1) the exact stage of differentiation of the cells (undifferentiated/uncommitted cells *versus* chondrogenic cells/chondrocytes); (2) the different nature of synchronisation applied (chemical/metabolic *versus* mechanical signals); (3) and the specific nature of the mechanical environment (stretch *versus* compression; cyclic *versus* static), which probably activate different downstream pathways that eventually result in clock entrainment. Nevertheless, the fact that properly set MS entrained the core clock of differentiating chondrocytes, and that it enhanced cartilage ECM production at least partially *via* synchronised cartilage marker gene and protein expression patterns compared to unstimulated cells is of key importance.

## 5. Conclusions and Perspectives

The circadian clock has been shown to play a key regulatory role in various tissues and organs, including the musculoskeletal system, and more specifically cartilage ^15,45^. Our results, for the first time, suggest that entraining chondroprogenitor clocks during chondrogenesis by uniaxial cyclic mechanical load offers a novel and insightful way in which these cells can be primed to produce more abundant cartilage ECM. Mechanical entrainment represents a non-invasive means by which the quality and quantity of tissue-engineered cartilage could be augmented through the control of the circadian clock in the critical period of chondrogenesis, avoiding the need for additional exogenous chemical or thermal stimuli. A deeper understanding of the biomechanical forces within the developing joint and their downstream signalling pathways *in vivo* may enhance our knowledge of postnatal articular cartilage development. Recapitulating the mechanical environment associated with cartilage formation and homeostasis during chondrogenic differentiation *in vitro* may promote the development of a cellular phenotype resembling chondrocytes in native articular cartilage, and advance the field of cartilage replacement strategies. It is therefore vital that a combined approach of a biomechanical environment and chondrochronology be considered when optimising future cartilage tissue engineering applications.

## Abbreviations

CCG: clock-controlled gene
CKI: casein kinase I
DMMB: dimethyl-methylene blue
DMSO: dimethyl sulfoxide
ECM: extracellular matrix
IHH: Indian hedgehog
LDS: longdaysin
MS: mechanical stimulation
MSC: mesenchymal stem cell
RORE: retinoic acid-related orphan receptor response element
RQ: relative quantity
RSD: relative standard deviation
TRPV: transient receptor potential vanilloid
TTFL: transcriptional-translational feedback loop

## Acknowledgements

The authors are thankful to Mrs. Krisztina Biróné Barna and Ms. Do Hwi Lee for technical assistance. We wish to thank Dr. Peter Nagy at the Department of Biophysics and Cell Biology, University of Debrecen, for developing the MATLAB-based image analysis software.

## Author Contributions

*Conception and design:* Csaba Matta, Róza Zákány; *Analysis and interpretation of the data:* Judit Vágó, Éva Katona, Roland Takács, Klaudia Dócs, Tibor Hajdú, Patrik Kovács, Daan van der Veen, Csaba Matta; *Drafting of the article:* Csaba Matta; *Critical revision of the article for important intellectual content:* all authors; *Final approval of the article:* all authors; *Statistical expertise:* Daan van der Veen; *Obtaining of funding:* Csaba Matta, Róza Zákány; *Administrative, technical, or logistic support:* Csaba Matta; *Collection and assembly of data:* Judit Vágó, Éva Katona, Roland Takács, Klaudia Dócs, Tibor Hajdú, Patrik Kovács.

