## Supporting information for "Cyclic uniaxial mechanical load enhances chondrogenesis through entraining the molecular circadian clock"

**This PDF file includes:**

- **Table S1.**	Forward and reverse primer sequences, gene accession numbers and
   amplicon length for each primer pair employed in this study
- **Table S2.**	Relative quantity (RQ) and relative SD (RSD) values for all genes analysed
   in this study after being normalised to the reference gene *RPL13*
- **Figure S1.**	Cell viability of chondrogenic micromass cultures following the loading scheme
   on day 6
- **Figure S2.**	Uncropped Simple Western Wes immunoassay electropherograms for the
   proteins analysed in this study
- **Figure S3.**	Global phosphorylation status of chondrifying micromass cultures following
   the loading regime, in combination with the clock modulator compound
   longdaysin

*1. Primer pairs for quantitative RT-PCR reactions*

**Table S1.** Forward and reverse primer sequences, gene accession numbers and amplicon length for each primer pair employed in this study

| **Gene** | **Accession number** | **Primer sequence** | **Product length** |
| --- | --- | --- | --- |
| *BMAL1* | NM_001001463.2 | CTGTGACAGAGGGAAGATACTGT | 119 |
|  |  | TGGCAATGTCTTTCGGATGAAGG |  |
| *PER2* | NM_204262.1 | CCAACTTTGTTGTGTGCTCCTTGC | 115 |
|  |  | CTGGAACAAACAGGTTGGTGTGTG |  |
| *PER3* | NM_001289779.1 | CGTTGGTACAGCCCTGTTTG | 180 |
|  |  | ACCCAGTTTTCAGTCGCTCAT |  |
| *CRY1* | NM_204245.1 | GGGTCTTCTTGCAACAGTGC | 143 |
|  |  | CGCCAACTGTCTGACCATCA |  |
| *CRY2* | NM_204244.1 | CAGGGGAGACCTCTGGATCA | 120 |
|  |  | AAGAATGCGCTGCACGAAAG |  |
| *REV-ERB* | NM_205205.2 | GAACACGTGACAGCTGGGTC | 120 |
|  |  | ATCACACCCCCTAAGGCAAG |  |
| *SOX6* | NM_001318451 | TGAATGGCTCTGCAGCTAAAC | 168 |
|  |  | TTCGACGCTCATCCTTAGCC |  |
| *SOX9* | NM_204281.1 | TTTCCGAGACGTGGACATCG | 150 |
|  |  | GTACCGCTGTAGGTGGTGAC |  |
| *RUNX2* | NM_204128 | GAGTCAGATTACAGACCCCAGG | 199 |
|  |  | AAATGGGCCCAGCTCGGAA |  |
| *ACAN* | NM_204955 | AGCAGTAGATGCACTGGGAC | 153 |
|  |  | GCCAGGTCGATCTCACACAG |  |
| *BGLAP* | NM_205387.4 | GGATGCTCGCAGTGCTAAAG | 147 |
|  |  | CTCAGCTCACACACCTCTCG |  |
| *COL1A2* | NM_001079714 | TATTGGAAGCCGTGGTCCATCT | 187 |
|  |  | ACCAGTGTCACCTCTCAGACC |  |
| *COL2A1* | NM_204426.1 | GGGACCTCAAGGCAAAGTCG | 140 |
|  |  | TTCCAGGCTCACCATTAGCG |  |
| *COL10A1* | NM_001396427.1 | TCCACAACATTTGAGGACGGA | 192 |
|  |  | TTCACCCCTCATCTGCACAC |  |
| *SPP1* | NM_204535 | CGAAGATCGCCACAGCATTG | 156 |
|  |  | GCCACGCGCTCTACATTTAC |  |
| *VEGF* | NM_205042 | CAGAGTCAGCACATAGCGCA | 169 |
|  |  | CACAGTGAAAGCTGGGTGGT |  |
| *RPL13* | NM_204999.1 | GGCCCGTGTTATCTCAGAGG | 113 |
|  |  | CCGCTTCTTTGGCACGTTTT |  |
| *YWHAZ* | NM_001031343 | GTTCCCTTGCAAAAACGGCT | 199 |
|  |  | GAGGCAGACGGAAGTTGGAA |  |
| *PPIA* | NM_001166326.1 | GAGCTCTTCGCTGACAAGGT | 139 |
|  |  | GCGTAAAGTCACCACCCTGA |  |

*2. Relative quantification data of normalised gene expression values*

**Table S2.** Relative quantity (RQ) and relative SD (RSD) values for all genes analysed in this study after being normalised to *RPL13* following mechanical load (**A**; upper table) or in non-stimulated chondrifying micromass cultures (**B**; lower table)

**A. Mechanical load**

| **Time points** |  | **24 h** | **32 h** | **40 h** | **48 h** | **56 h** | **64 h** | **72 h** |
| --- | --- | --- | --- | --- | --- | --- | --- | --- |
| *BMAL1* | RQ | 1.000 | 1.109 | 0.842 | 0.856 | 0.741 | 0.404 | 0.374 |
|  | RSD | 0.057 | 0.115 | 0.072 | 0.340 | 0.056 | 0.031 | 0.109 |
| *CRY1* | RQ | 1.000 | 1.006 | 0.884 | 0.809 | 0.937 | 0.444 | 0.529 |
|  | RSD | 0.058 | 0.195 | 0.086 | 0.011 | 0.223 | 0.157 | 0.116 |
| *CRY2* | RQ | 1.000 | 0.835 | 0.903 | 0.887 | 0.901 | 0.475 | 0.512 |
|  | RSD | 0.085 | 0.181 | 0.195 | 0.119 | 0.153 | 0.178 | 0.189 |
| *PER2* | RQ | 1.000 | 1.174 | 0.635 | 0.643 | 0.696 | 0.407 | 0.551 |
|  | RSD | 0.092 | 0.029 | 0.045 | 0.083 | 0.119 | 0.086 | 0.027 |
| *PER3* | RQ | 1.000 | 1.038 | 0.737 | 0.913 | 1.001 | 0.375 | 0.644 |
|  | RSD | 0.360 | 0.170 | 0.152 | 0.260 | 0.244 | 0.209 | 0.089 |
| *REV-ERB* | RQ | 1.000 | 2.692 | 1.200 | 1.070 | 1.060 | 1.650 | 2.692 |
|  | RSD | 0.231 | 0.241 | 0.028 | 0.125 | 0.057 | 0.050 | 0.156 |
| *SOX9* | RQ | 1.000 | 4.240 | 0.911 | 1.540 | 1.215 | 0.643 | 0.620 |
|  | RSD | 0.195 | 0.083 | 0.018 | 0.165 | 0.092 | 0.037 | 0.037 |
| *COL2A1* | RQ | 1.000 | 1.111 | 0.685 | 1.591 | 0.992 | 0.974 | 0.753 |
|  | RSD | 0.113 | 0.248 | 0.210 | 0.128 | 0.097 | 0.029 | 0.107 |
| *ACAN* | RQ | 1.000 | 1.945 | 1.280 | 1.454 | 1.265 | 0.610 | 0.748 |
|  | RSD | 0.021 | 0.048 | 0.073 | 0.019 | 0.051 | 0.062 | 0.040 |
| *SOX6* | RQ | 1.000 | 2.583 | 1.022 | 0.902 | 0.632 | 0.307 | 0.451 |
|  | RSD | 0.034 | 0.054 | 0.020 | 0.003 | 0.067 | 0.140 | 0.058 |
| *RUNX2* | RQ | 1.000 | 1.638 | 1.390 | 1.112 | 0.654 | 0.533 | 0.567 |
|  | RSD | 0.289 | 0.235 | 0.093 | 0.111 | 0.259 | 0.112 | 0.095 |

**B. Non-stimulated (control) cultures**

| **Time points** |  | **24 h** | **32 h** | **40 h** | **48 h** | **56 h** | **64 h** | **72 h** |
| --- | --- | --- | --- | --- | --- | --- | --- | --- |
| *BMAL1* | RQ | 1.000 | 1.013 | 0.993 | 0.830 | 0.850 | 0.977 | 0.938 |
|  | RSD | 0.136 | 0.163 | 0.093 | 0.104 | 0.206 | 0.137 | 0.134 |
| *CRY1* | RQ | 1.000 | 1.100 | 1.235 | 1.134 | 1.242 | 1.168 | 1.133 |
|  | RSD | 0.150 | 0.071 | 0.035 | 0.047 | 0.100 | 0.090 | 0.070 |
| *CRY2* | RQ | 1.000 | 1.066 | 1.070 | 1.071 | 1.000 | 1.026 | 1.055 |
|  | RSD | 0.096 | 0.063 | 0.031 | 0.091 | 0.055 | 0.058 | 0.047 |
| *PER2* | RQ | 1.000 | 0.846 | 0.829 | 0.867 | 1.013 | 0.940 | 0.883 |
|  | RSD | 0.086 | 0.081 | 0.118 | 0.154 | 0.076 | 0.077 | 0.103 |
| *PER3* | RQ | 1.000 | 0.921 | 0.965 | 0.918 | 0.860 | 0.940 | 0.926 |
|  | RSD | 0.250 | 0.027 | 0.068 | 0.052 | 0.089 | 0.130 | 0.069 |
| *REV_ERB* | RQ | 1.000 | 1.029 | 1.085 | 1.209 | 1.120 | 1.006 | 1.077 |
|  | RSD | 0.135 | 0.104 | 0.129 | 0.097 | 0.078 | 0.099 | 0.085 |
| *SOX9* | RQ | 1.000 | 0.791 | 0.939 | 0.961 | 1.038 | 0.995 | 0.837 |
|  | RSD | 0.151 | 0.061 | 0.080 | 0.103 | 0.096 | 0.043 | 0.051 |
| *COL2A1* | RQ | 1.000 | 0.972 | 0.885 | 0.827 | 0.893 | 0.994 | 0.929 |
|  | RSD | 0.082 | 0.057 | 0.053 | 0.043 | 0.052 | 0.066 | 0.045 |
| *ACAN* | RQ | 1.000 | 0.917 | 0.953 | 0.951 | 1.149 | 1.211 | 0.924 |
|  | RSD | 0.079 | 0.007 | 0.084 | 0.067 | 0.054 | 0.066 | 0.039 |
| *SOX6* | RQ | 1.000 | 2.872 | 2.722 | 3.126 | 2.407 | 2.800 | 3.418 |
|  | RSD | 0.036 | 0.077 | 0.148 | 0.120 | 0.104 | 0.069 | 0.098 |
| *RUNX2* | RQ | 1.000 | 0.864 | 0.880 | 0.893 | 0.981 | 1.034 | 1.007 |
|  | RSD | 0.015 | 0.045 | 0.099 | 0.035 | 0.055 | 0.063 | 0.052 |

*3. Cell viability (MTT assay)*

Cell viability was monitored using the MTT assay. Micromass colonies were seeded into 6-well plates and the loading regime was carried out as with other cultures (i.e., 60 min daily for 6 days). Immediately after the last loading cycle on day 6, 50 µL of MTT reagent (3-[4,5-dimethyl-2-thiazolyl]-2,5-diphenyl-2H-tetrazolium bromide; 5 mg/mL in PBS; Sigma-Aldrich, Merck; Cat. no: M5655) were pipetted into each well. Cells were incubated for 2 hours at 37 °C. Following the addition of 500 μL MTT solubilizing solution (10% Triton X-100 in 2-propanol), optical density was measured at 570 nm (Chameleon, Hidex Ltd., Turku, Finland). Measurements were carried out in 3 samples of each experimental group in 3 independent experiments. Optical density readings of the experimental group have been normalized to those of the controls and shown as percentage changes (*Figure S1*). The data indicate that the applied loading regime did not influence mitochondrial activity/cell survival/cell death rates in chondrifying micromass cultures.

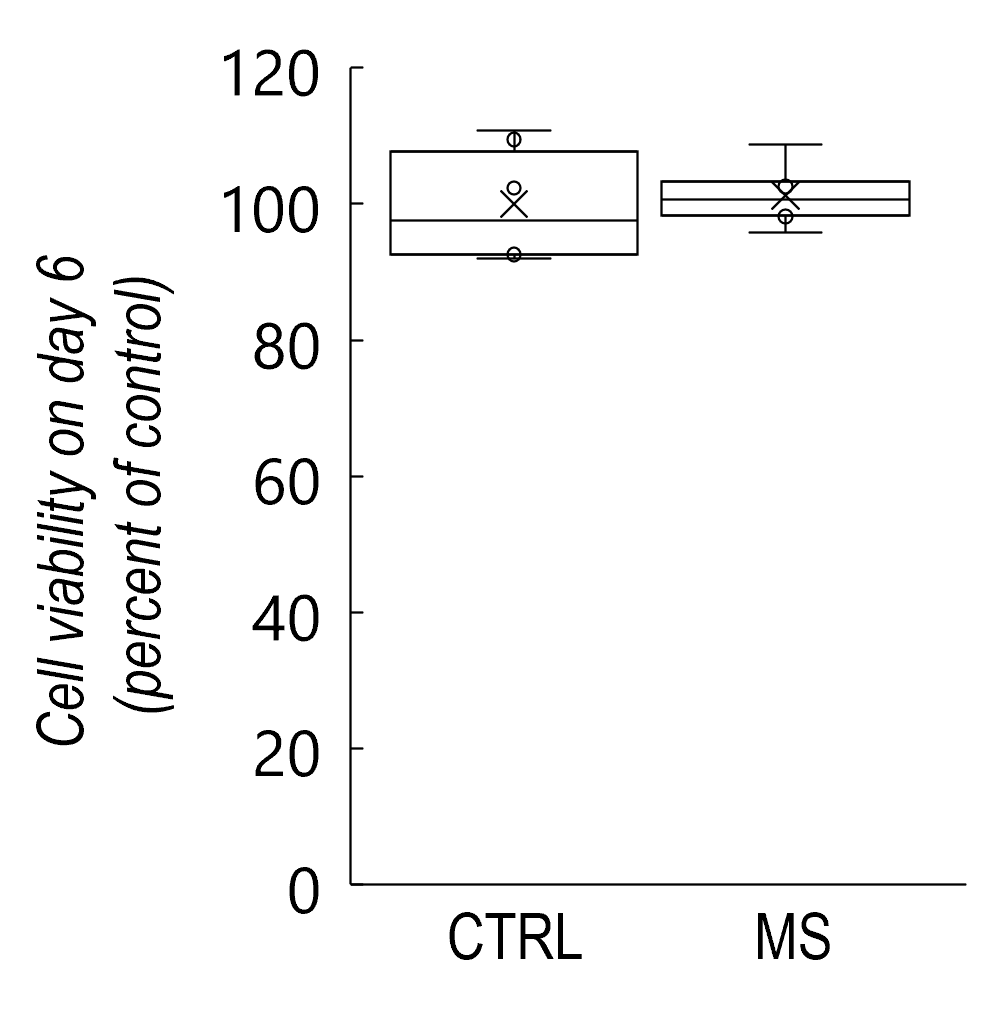

*Figure S1.* Cell viability of 6-day-old micromass cultures following the mechanical loading regime *versus* the untreated control cultures, as determined by MTT assay (representative data).

*4. Uncropped Simple Western Wes electropherograms*

*Figure 6A* in the manuscript shows Simple Western Wes electropherograms at cropped the specific/expected molecular weight for each primary antibody. Below are the uncropped electropherograms for the primary antibodies shown in *Table 3*.

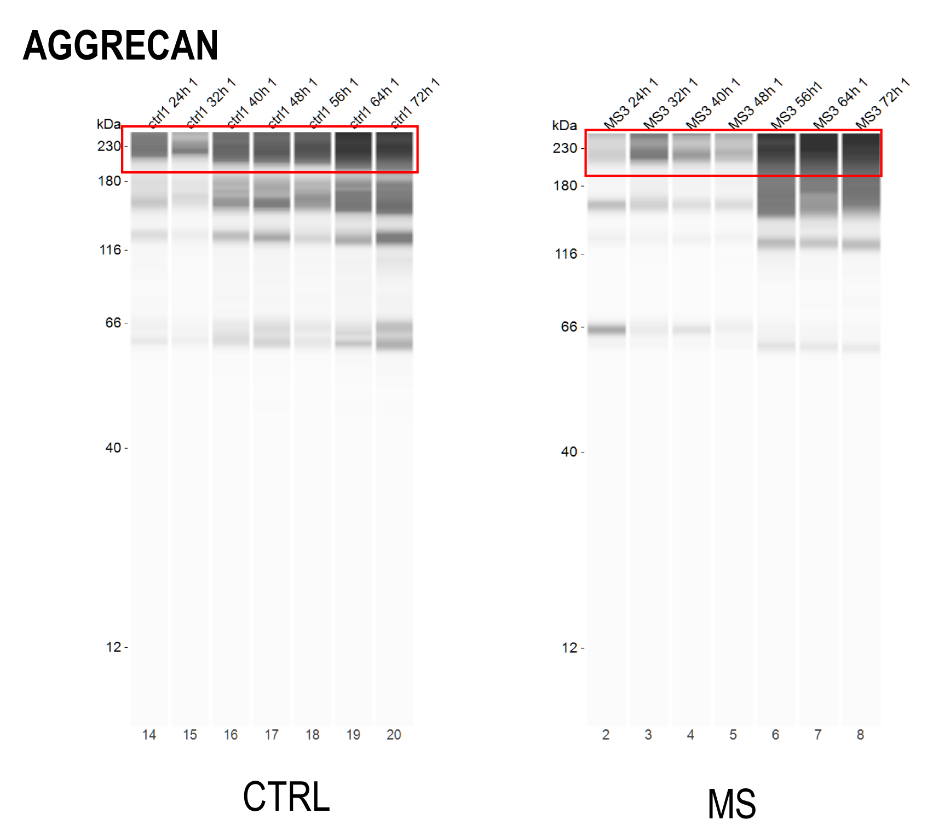

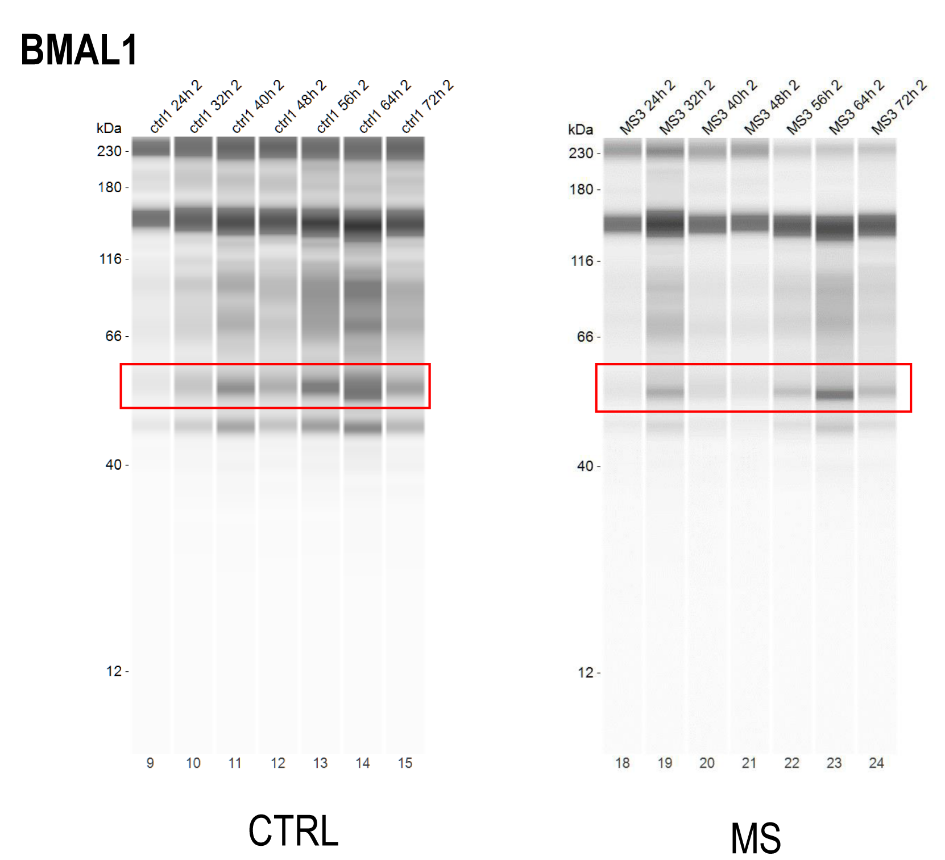

*Figure S2 (continued)*

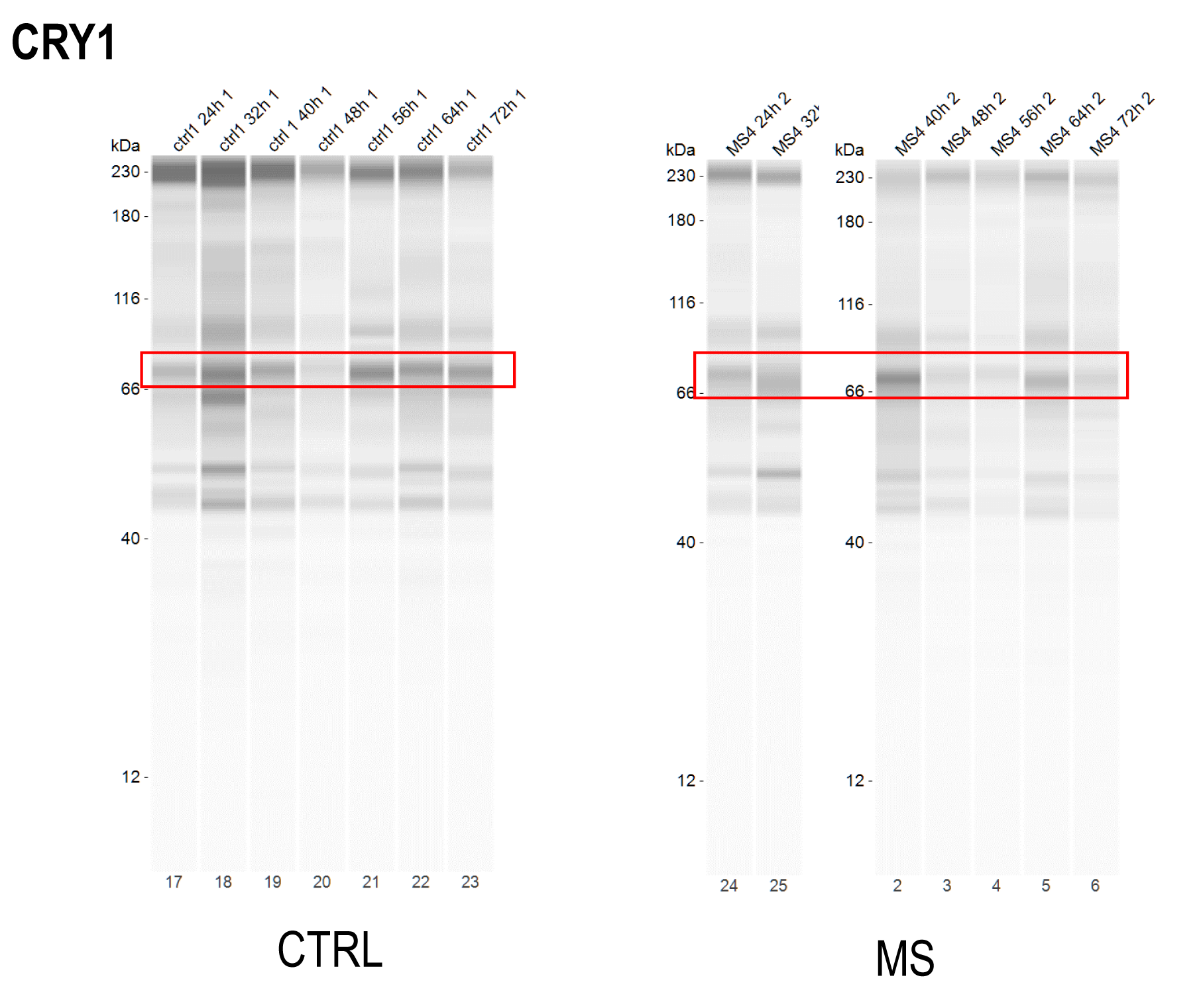

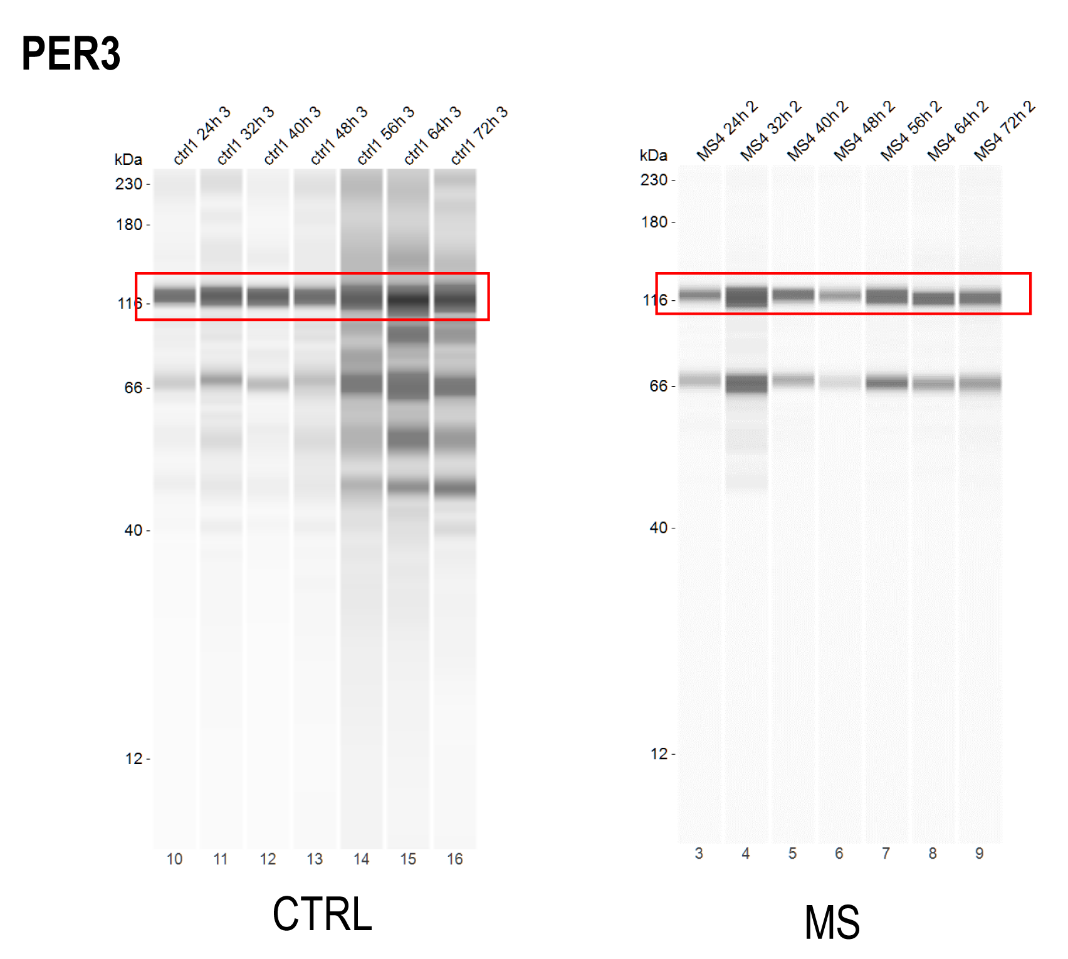

*Figure S2 (continued)*

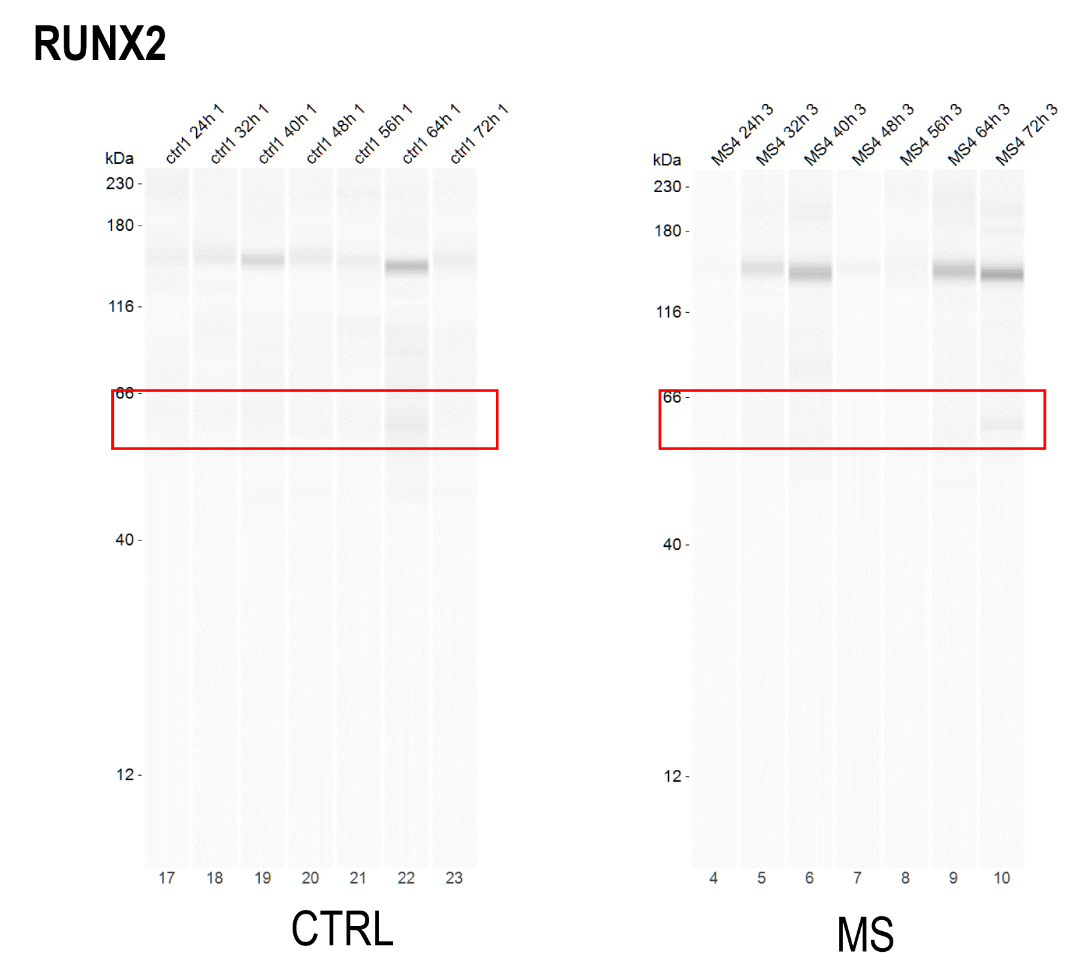

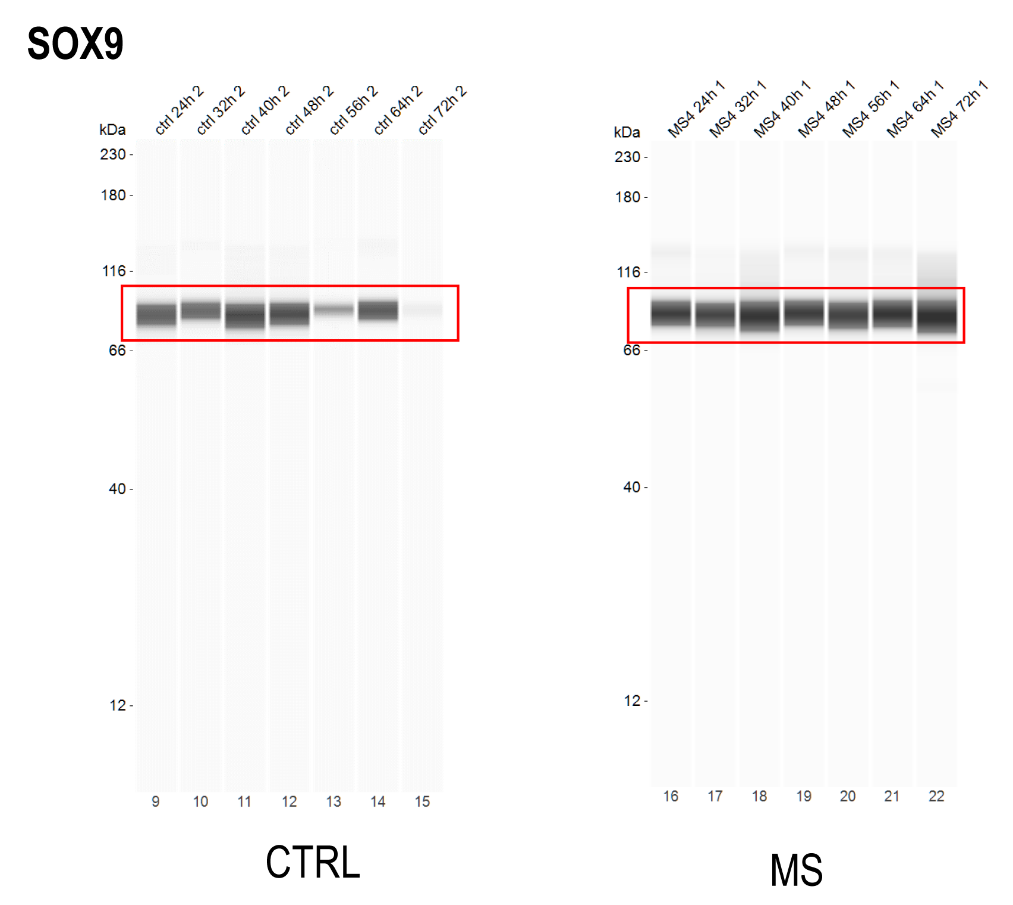

*Figure S2 (continued)*

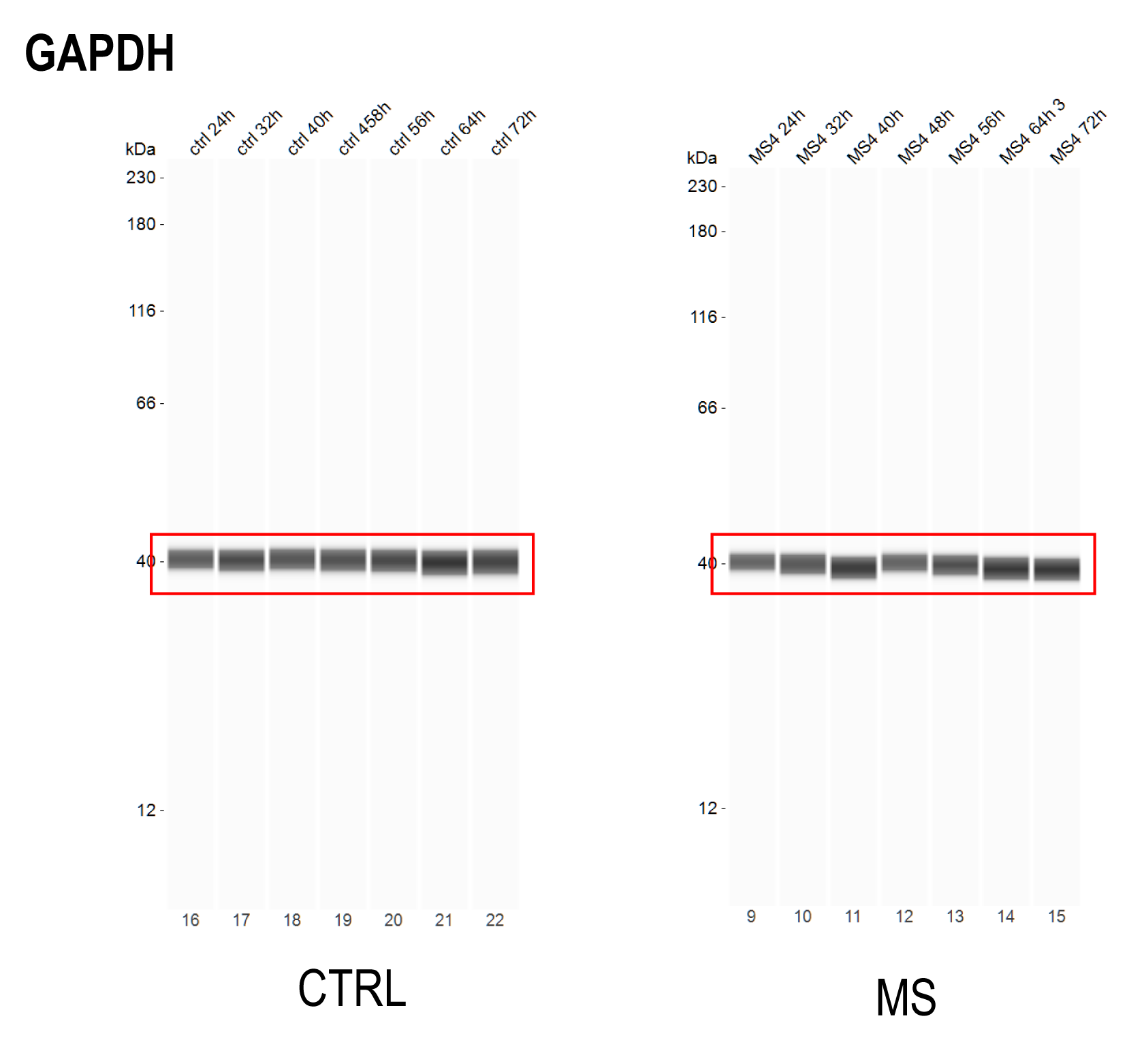

*Figure S2.* Uncropped Simple Western Wes electropherograms for the cropped bands at expected molecular weights presented in *Figure 6*.

*5. Phosphorylation status in micromass cultures following the loading scheme with or without
5 µM longdaysin*

To check whether the mechanical loading scheme and/or application of the clock modulator compound longdaysin at 5 μM for 60 minutes each day exerted detectable changes at the global phosphorylation status in chondrifying micromass cultures, we performed conventional western blots on total lysates and probed with a pan phospho Serine/Threonine primary antibody.

**Materials & Methods**

**SDS-PAGE and western blot analysis**

Micromasses following the last mechanical loading scheme and/or the last LDS challenge on day 6 were washed in PBS and then harvested. After centrifugation, cell pellets were suspended in 100 μL of RIPA (Radio Immuno Precipitation Assay) homogenization buffer (composed of 150 mM NaCl; 1.0% NP40, 0.5% sodium deoxycholate; 50 mM Tris, 0.1% SDS; pH 8.0) supplemented with protein inhibitors as follows: aprotinin (10 µg/mL), 5 mM benzamidine, leupeptin (10 µg/mL), trypsin inhibitor (10 µg/mL), 1 mM PMSF, 5 mM EDTA, 1 mM EGTA, 8 mM Na-fluoride, 1 mM Na-orthovanadate. All components were purchased from Sigma-Aldrich. Samples were stored at −80°C.

Suspensions were sonicated by pulsing burst for 30 sec, at 40 A by 50 cycles using an ultrasonic homogeniser (Cole-Parmer, Vernon Hills, IL, USA). As total cell lysates were used for western blotting, samples for sodium dodecyl sulphate-polyacrylamide gel electrophoresis (SDS-PAGE) were prepared by the addition of Laemmli’s electrophoresis sample buffer (4% SDS, 10% ß-mercaptoethanol, 20% glycerol, 0.004% bromophenol blue, 0.125 M Tris–HCl; pH 6.8) to cell lysates to set equal protein concentration (0.5 µg/mL), and boiled for 10 min at 95°C. 20 µg of protein was separated by 10% SDS-PAGE for the detection of serine/threonine phosphorylated proteins and actin. Separated proteins were transferred to nitrocellulose membranes (Bio-Rad Trans Blot Turbo Midi Nitrocellulose Transfer Packs; Bio-Rad Laboratories, Hercules, CA, USA) by using a Bio-Rad Trans-Blot Turbo system. After blocking in 5% non-fat dried milk (PanReac AppliChem ITW Reagents, Darmstadt, Germany) in PBS for 1 h, membranes were washed and exposed to primary antibodies overnight at 4°C in the dilution as given in *Table S3* below. After washing for 30 min in PBST (PBS supplemented with 1% Tween-20 [Amresco Inc., Solon, OH, USA]), membranes were incubated with Alexa Fluor 647 conjugated anti-rabbit IgG secondary antibodies (Life Technologies Corporation, Carlsbad, CA, USA) in 1:1000 dilution. Fluorescent signals were developed with a gel imaging system (ChemiDoc MP Imaging System, Bio-Rad Laboratories).

| ***Antibody*** | *Host animal* | *Dilution* | *Vendor, Cat. no.* |
| --- | --- | --- | --- |
| **Anti-pan Phospho Serine/Threonine** | rabbit, polyclonal | 1:100 | Sigma-Aldrich; SAB5700525 |
| **Anti-ß-Actin** | mouse, monoclonal | 1:5000 | Sigma-Aldrich; A5441 |

*Table S3.* Table of antibodies used in the experiments

We could not detect marked changes in the global phosphorylation status in micromass cultures either following the loading scheme or following treatment with LDS with this method (*Figure S3*).

**A.**

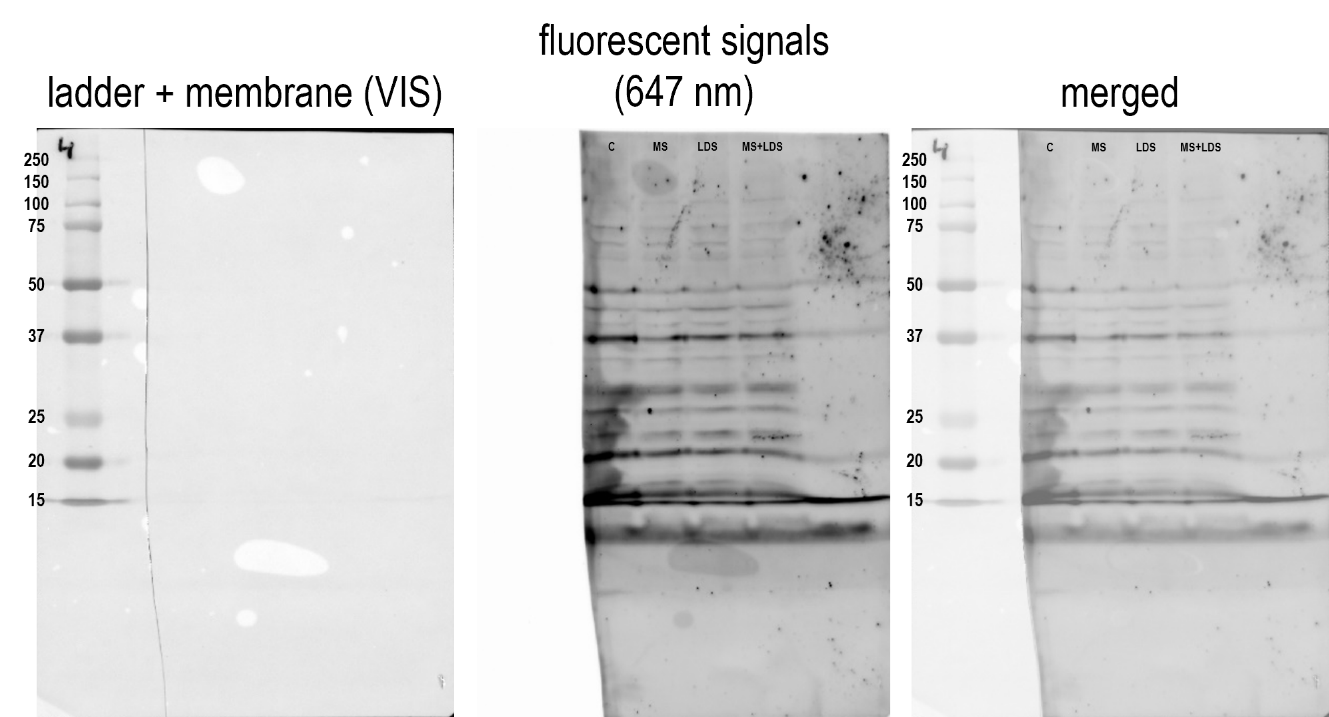

**B.**

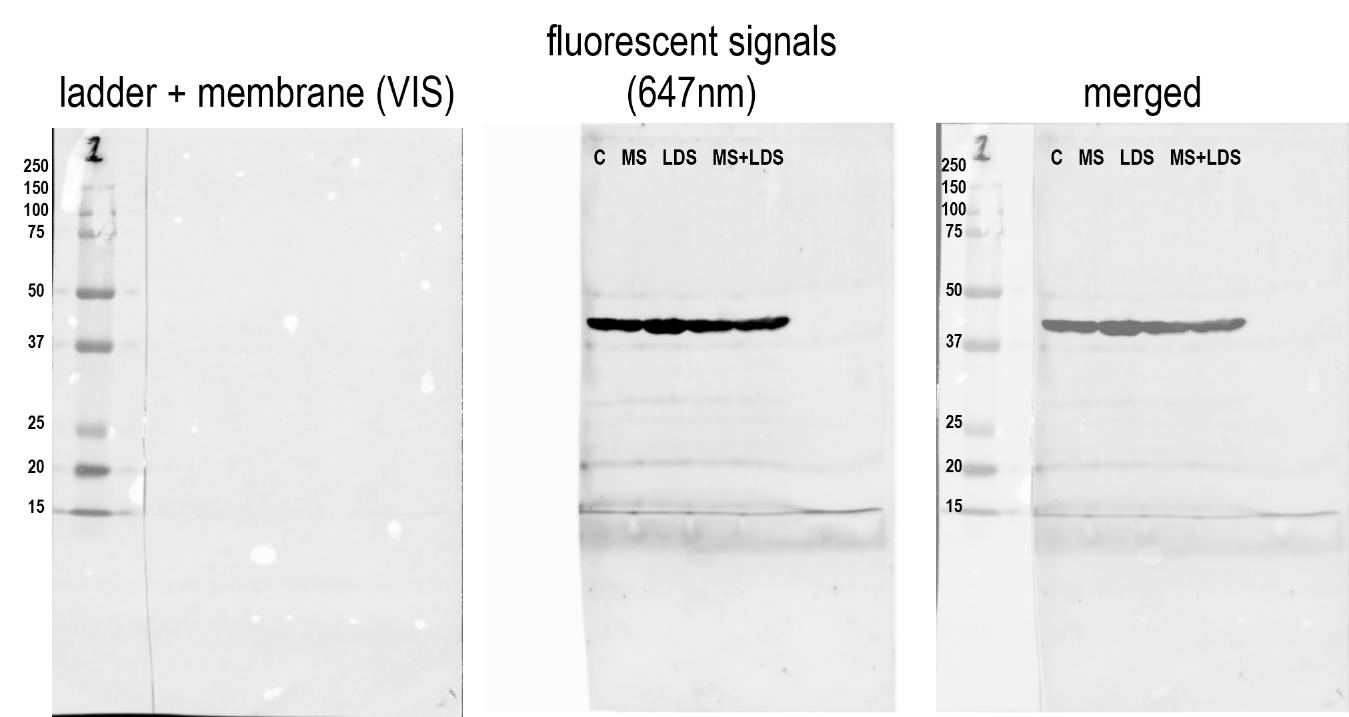

*Figure S3.* Conventional western blot results performed on total lysates. **A.** Membrane probed with the anti-pan phosphor serine/threonine primary antibody. **B.** Membrane probed with anti-actin primary antibody.
